# A new family of ribosome hibernation factors in Archaea

**DOI:** 10.1101/2025.10.11.676729

**Authors:** Clément Madru, Gabrielle Bourgeois, Rémi Dulermo, Régine Capeyrou, Karima Figuigui, Claire Duboc, Stéphane l’Haridon, Logan McTeer, Marta Kwapisz, Béatrice Clouet-d’Orval, Marie Bouvier, Yves Mechulam, Guillaume Borrel, Emmanuelle Schmitt, Didier Flament

## Abstract

Ribosome hibernation preserves translation machinery during stress, yet its mechanisms in Archaea remain poorly defined. Here we identify Hib, a previously unrecognized family of hibernation factors widespread in Archaea. Deletion of *hib* in *Thermococcus barophilus* delays recovery from stationary phase and reduces 70S ribosome pools, establishing its role in ribosome preservation. Hib displays a unique modular architecture, combining a bacterial-like HPF/RaiA domain with a Cystathionine Beta Synthase module. High-resolution cryo-EM structures of reconstituted and in cell-extracted Hib:ribosome complexes from *Pyrococcus abyssi* identify three conformations encompassing the positions of tRNAs at A, P and E sites during translation. Thus, Hib acts as a hibernation factor blocking all states of the dynamic translation process, preserving the ribosome from dissociation and degradation. Our findings define Hib as a key hibernation factor in Archaea and provide a framework for understanding ribosome dormancy and adaptation across all domains of life. Beyond the discovery of Hib, a comprehensive phylogenetic analysis highlights the evolutionary relationships between prevalent ribosome hibernation factors across Bacteria and Archaea.

## Introduction

Protein synthesis is an essential and highly energy-demanding process that requires tight regulation, particularly under stress or nutrient limitation^1,2^. A major strategy to modulate translation in these conditions is ribosome hibernation, a mechanism that allows cells to reversibly inactivate and stabilize ribosomes, thereby conserving resources while preserving the translational machinery for rapid reactivation once favourable conditions return^3–6^. In Bacteria and Eukaryotes, ribosome hibernation is mediated by dedicated protein factors that block the binding of mRNA and tRNA, halting translation and protecting ribosomes from degradation of their active centres by cellular nucleases^7,8^.

Despite their structural diversity, the loss of these factors contributes to similar phenotypes, i.e. reduced growth resumption, reduced stress tolerance and accelerated ribosome degradation^9–12^. Bacterial hibernating ribosomes can exist as a translationally inactive 70S particles or dimerized 100S complexes, stabilized by protein factors classified in three different families; HPF/RaiA, RMF and Balon^13–16^. HPF/RaiA proteins (lHPH, HPF & RaiA) share a structural domain that binds the 30S small ribosomal subunit (SSU) on the binding sites of mRNA-, tRNA- and initiation factors to inactivate the ribosome^17^. In contrast, Balon binds to ribosomes in an mRNA-independent manner, thereby enabling the initiation of hibernation even while ribosomes are engaged in protein synthesis^14^. Finally, RMF induces conformational changes in the 30S subunit, promoting the formation of 100S dimers^18^. Eukaryotic ribosome hibernation factors are more diverse and include six families: Stm1/Serbp1, Lso2, IFRD1/IFRD2, MDF1, MDF2 and Dap1b (reviewed in^14^). These factors were shown to act in a similar way to bacterial factors, by occupying the mRNA channel and either partially or fully blocking tRNA binding sites. However, eukaryotic and bacterial hibernation factors are not homologous indicating an independent origin^5^.

By contrast, ribosome hibernation in Archaea remains largely unexplored, despite its likely importance for survival in extreme and nutrient-limited environments. The archaeal ribosome is of the eukaryotic type. With the exception of the 3’ end of the 16S rRNA, rRNAs of archaeal ribosomes are closer to eukaryotic rRNAs than to bacterial rRNAs^19–23^. In addition, Archaeal ribosomal proteins are either universal or found only in Eukaryotes and Archaea^22–26^. To date, only two archaeal ribosome hibernation factors have been recently reported. aRDF, restricted to the *Pyrococcus* genus, promotes small-subunit dimerization by binding to the ribosomal protein eS32 thereby preventing 70S assembly through a unique head-to-body architecture^27^. A second hibernation factor, named Dri, has been recently identified in *Pyrobaculum calidifontis*. Dri is a dual-lobed Cystathionine Beta Synthase (CBS) modules protein that binds the 30S and 50S subunits, blocking the mRNA channel and the peptidyl transferase centre, thereby inactivating translation^28^. Its function as a hibernation factor is supported by biochemical and structural analyses but no genetic evidence for its physiological role in the cells is available. Furthermore, Dri homologues are unevenly distributed, being largely restricted to *Thermoproteota* or present as single CBS modules, in a subset of other phyla.

Here, we report the functional and structural characterization of a novel family of archaeal ribosome hibernation factors, which we have designated Hib. Combining genetic, phylogenomic, biochemical and structural approaches, we demonstrate that Hib associates with both ribosomal subunits to inactivate ribosomes by blocking the mRNA channel and tRNA binding sites. Deletion of *hib* in the model archaeon *Thermococcus barophilus* results in a significant delay in growth resumption from stationary phase, as well as a marked reduction in 70S ribosome particles. This underscores the significance of ribosome stabilization for physiological processes during stress periods. High-resolution cryo-EM structures of reconstituted and cell-extracted Hib:ribosome complexes from *Pyrococcus abyssi* reveal three conformations encompassing the positions of tRNAs at A, P and E sites during translation. Thus, Hib acts as a hibernation factor blocking all states of the dynamic translation process. In cell-extracted Hib:ribosome complexes, we identify an E-site tRNA, known to act as an aa-tRNA deprivation sensor, as a partner of Hib. Furthermore, we propose that archaeal homologue of SBDS (Shwachman-Bodian-Diamond Syndrome) is involved in Hib-mediated hibernation. Finally, we observed hibernating ribosomes containing a Shine-Dalgarno (SD):antiSD duplex. Thus, the preservation of ribosomes in *Thermococcales* archaea is achieved through the protection of all active sites, from the mRNA entry channel to the mRNA exit channel.

Together, our findings establish Hib as a key mediator of ribosome hibernation in Archaea. They provide a mechanistic and functional framework to understand how archaeal cells preserve their translational machinery under stress, and open new perspectives on the evolution and regulation of ribosome dormancy across the three domains of life.

## Results

### Identification of Hib as a novel, widely distributed hibernation factor in Archaea

To identify potential archaeal hibernation factors, we performed PSI-Blast searches against archaeal genomes using bacterial and eukaryotic ribosome hibernation factors as seeds. This search revealed a large protein family with low sequence identity with bacterial HPF/RaiA proteins. These proteins were detected in 47% of the archaeal genomes and are widely distributed across all major lineages of Archaea. Members of this family, typically annotated as Cystathionine Beta Synthase (CBS) domain-containing proteins, display a unique modular architecture. The N-terminal part is composed of four CBS domains of about 60 amino acids, arranged in tandem (hereafter named CBS module or CM), linked to a C-terminal part that shares sequence similarities with the core domain of bacterial hibernation promoting factors (Supplementary Fig. 1). We propose to name this new protein family Hib. The comparative alignment of representative members of the bacterial RaiA, HPF, and lHPF families with Hib proteins from diverse archaeal phyla showed conservation of secondary structural elements in the C-terminal domain. In addition, we observed two insertions specific of archaeal Hib proteins (Supplementary Fig. 1).

We next tested whether Hib associates with the ribosomal subunits. We produced and purified a His-tagged recombinant version of Hib from *Pyrococcus abyssi* (Pa-Hib, uniprot: Q9UYR4) and performed gel filtration analysis experiments with purified *P. abyssi* ribosomal subunits. As shown in Fig. 1a and Supplementary Fig. 2, Hib consistently co-eluted with either a mixture of 30S and 50S or with 30S alone, indicating that Hib interacts strongly with the SSU alone or in the context of the full ribosome.

**Fig. 1:**
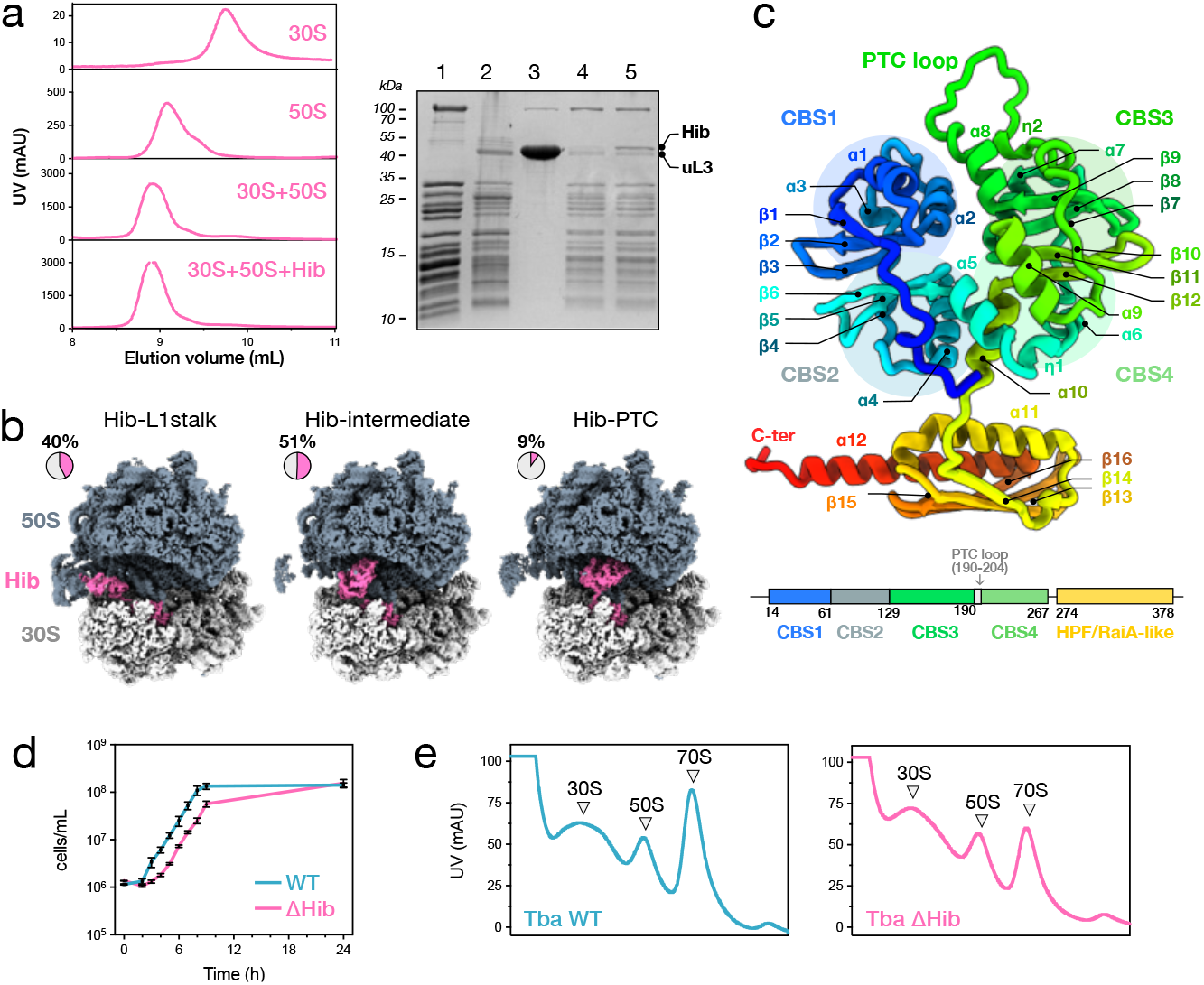
Hib is a hibernation factor. a. Bio-Agilent Sec5 chromatograms for *P. abyssi* ribosomal subunits, 30S, 50S, 30S+50S mixture, 30S+50S+Hib mixture). SDS-PAGE analysis is shown in the right-hand corner. 1:30S, 2:50S, 3:Hib, 4:elution peak of the 30S+50S mixture, 5: elution peak of the 30S+50S+Hib mixture. The analysis shows that Hib coelutes with the 70S fraction. The high molecular protein corresponds to phosphoenol pyruvate synthase (PEP) that is a usual contaminant of the 30S preparation. b. Cryo-EM structures of reconstituted Pa-70S:Hib complex at 2.5 Å resolution. Data processing and 3D classifications identified three conformational states distinguished by the position of the Hib N-terminal domain. c. Hib 3D structure. The N-terminal domain of Hib (1-269) is made up of four CBS domains. Two CBS domains interacts through their beta-sheets to form a bateman module. Two bateman modules interact to form a disk-like structure defined as parallel CBS module in^51^. Each CBS domain consists of three-standed (two parallel and one anti-parallel) β-sheet with two α-helices^52^. In domains 2,3 and four the first beta strand is less well defined and extended, as already mentionned in other CBS-domain containing structures. One long loop is inserted between α8 and β10 of the third CBS domain. The C-terminal domain (272-387) has the βαβββα topology found in all bacterial HPF/RaiA-like structures. The five last residues are not visible in the cryo-EM map. d. Δ517 (blue) and Δ*hib* (pink) cells grown in TRM medium at 85°C for 24h, were used for inoculating fresh TRM medium. Cells were then incubated at 85°C and their growth was monitored by flow cytometry at various time after inoculation. e. “3 mg of total proteins, from Δ517 (blue) and Δ*hib* (pink) cells in stationary phase (15h), were fractionated on 10 to 50% sucrose density gradients by ultracentrifugation. The free 30S and 50S ribosomal subunits and 70S ribosomes are indicated. The polysomes fractions were cropped.

To confirm these results, we reconstituted 70S:Hib complexes *in vitro* for cryo-EM analyses. We collected a large data set (∼30k movies) on a Titan Krios microscope (ExtendedData Fig. 1) and obtained ∼840k 70S particles. 3D classification identified three distinct 70S:Hib conformations (Fig. 1b). In all three classes, the C-terminal domain is located in the active sites (A, P, E) of the SSU, in a position that closely resembles that of HPF homologues^14,17,18,29^. In contrast, the N-terminal domain adopted three different positions on the large ribosomal subunit (LSU): bound to the peptidyl-transferase site (PTC) (Hib-PTC; 9% of the Hib-bound particles), between P and E sites (Hib-intermediate; 51% of the Hib-bound particles), or at the E site contacting the L1-stalk (Hib-L1Stalk; 40% of the Hib-bound particles). Comparison of Hib-PTC and Hib-intermediate cryo-EM maps showed that the ribosome is in the same unrotated state. In contrast, Hib-L1Stalk corresponds to a rotated state of the 30S with respect to the 50S. In addition to 70S ribosomes, we also identified Hib-bound 30S particles (ExtendedData Fig. 1). In these ribosomal complexes, the C-terminal domain occupied the mRNA binding channel, while the N-terminal domain remained unresolved, consistent with increased mobility in the absence of the LSU. Together, these data show that Hib occupies and blocks ribosomal active sites on both ribosomal subunits, displaying characteristics similar to those of bacterial and eukaryotic hibernation factors.

Hib was first built into the 2.9 Å cryo-EM map obtained from the Hib-PTC conformation, stating from an AlphaFold model. Pa-Hib is a 392–amino acid protein with a bipartite architecture: an N-terminal domain, residues 1–269, containing four CBS domains arranged in tandem and forming a CBS module, connected by a short linker to a C-terminal domain (residues 272– 392) that adopts the βαββα fold characteristic of bacterial HPF/RaiA proteins (Fig. 1c).

To further investigate the function of Hib, we generated a *hib*-deleted mutant strain of *Thermococcus barophilus* (uniprot: F0LKU5), a genetically tractable *Thermococcales* species from the same ecological niche as *P. abyssi*^30^. As shown in Fig. 1d, regrowth of stationary-phase Δ*hib* mutant cells was significantly delayed compared with that of wild-type cells: lag phases for WT and Δ*hib* lasted 2 and 4h, respectively. To test whether this phenotype reflected reduced viability of the Δ*hib* mutant, we performed Most Probable Number assays after culturing the cells in nutriment-deprived medium for up to 120h. Viable cell counts were comparable across WT and Δ*hib* (Supplementary Fig. 3), indicating that the growth delay is not directly related to cell viability under the tested conditions. Instead, sucrose gradient analyses revealed that the number of 70S ribosomes in Δ*hib* stationary-phase cells was ∼60% lower than in WT, as determined by peak area comparisons (Fig. 1e). This substantial decrease in 70S particles demonstrates that Hib is crucial for maintaining the stability and integrity of the inactive ribosome pool during multi-stress stationary phase. It also suggests that the impaired growth recovery of the Δ*hib* strain likely stems from the time needed to rebuild a sufficient pool of functional ribosomes before translation and growth can restart.

### Cryo-EM structures from cell extracts reveal multiple Hib:ribosome conformations

To strengthen our functional analyses, we isolated Hib-bound ribosomes directly from *P. abyssi* cell extracts^31^. Instead of using a standard ribosome purification protocol, we only performed one sucrose gradient step to recover the high-molecular weight fraction which was directly used for a large cryo-EM data collection. 70S particles were easily picked from the images. In addition, we also detected many macromolecular complexes, the description of which goes beyond the scope of this article (Fig. 2a-b and ExtendedData Fig. 2). Overall, the large variety of high-molecular weight complexes observed in the images validates our purification strategy aimed at isolating states close to the native state.

**Fig. 2:**
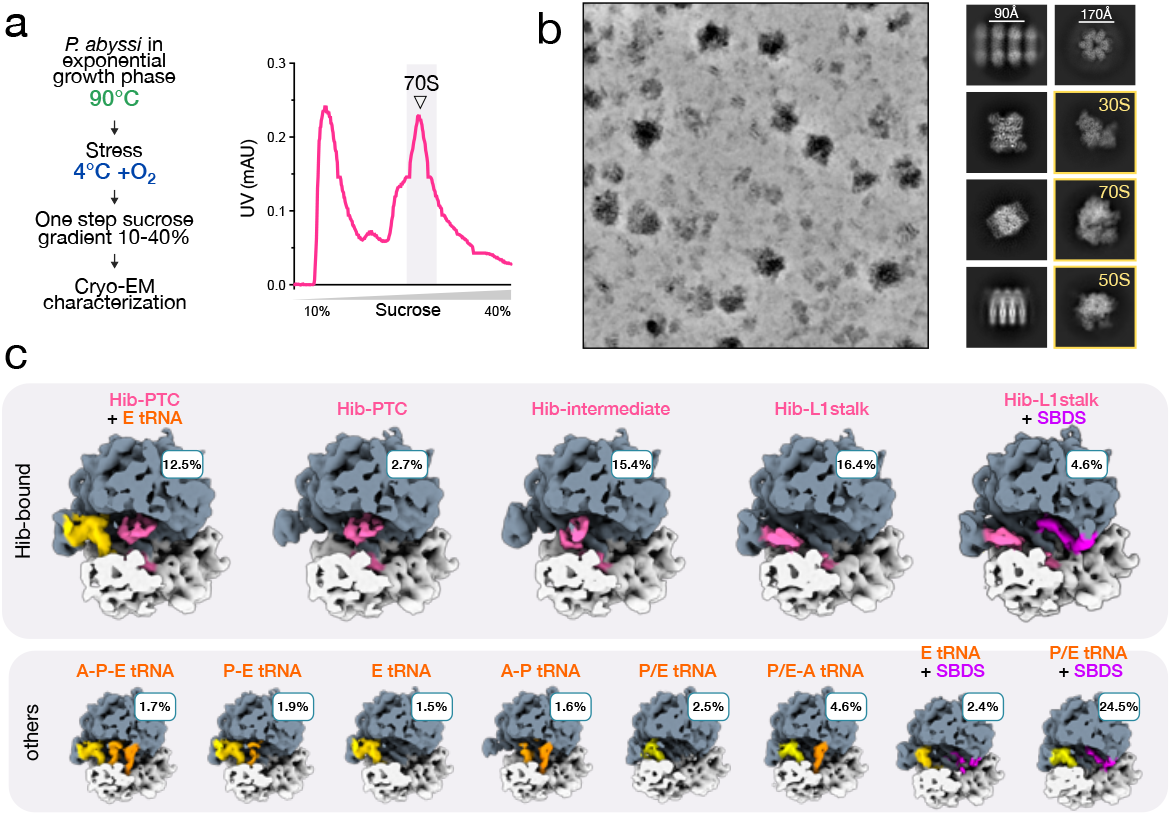
Cryo-EM structures reveal multiple Hib:ribosome complexes in cell extracts. a. Schematic overview of the experimental protocol. *P. abyssi* cells were grown at 90 °C (exponential phase), exposed to cold stress at 4 °C for 1 h, and subjected to oxidative stress during cell harvesting. Ribosomal complexes were isolated by one-step sucrose gradient centrifugation, and the 70S fraction was analyzed by cryo-EM. The corresponding sucrose gradient profile is shown. b. Representative denoised micrograph (processed with cryoSPARC) illustrating the presence of high-molecular-weight complexes, together with corresponding 2D classes. c. 3D classes obtained from the 70S particle dataset, with the relative proportion of each class indicated.

We isolated ∼1.5 M 70S particles, 51.6% of which contained Hib. 3D classification showed that ribosome:Hib complexes adopted the same three conformations previously observed in the reconstituted complexes (Fig.1b and 2c). However, this time, Hib-PTC accounts for ∼30 % of the Hib containing particles. Also, most of the Hib-PTC particles contained an E site tRNA (Fig. 2c), suggesting that it promotes the stabilization of the Hib-PTC conformation. In addition, an unexpected class was found to have both Hib and SBDS bound to the ribosome (Fig. 2c).

Heterogeneity and conformational variability were addressed using 3D flex refinement in CryoSparc (ExtendedData Fig. 2).

In all conformational states, the Hib C-terminal domain occupies the same position within the mRNA:tRNA binding channel. This domain establishes extensive contacts with the ribosome (Fig. 3c, e-f). In the mRNA binding channel, Hib-α12 helix contacts h44, h28, h24, h23 helices of the 16S rRNA as well as uS7, uS11 and eS28 ribosomal protein (Supplementary Fig. 1 and SupplementaryData 1).

**Fig. 3:**
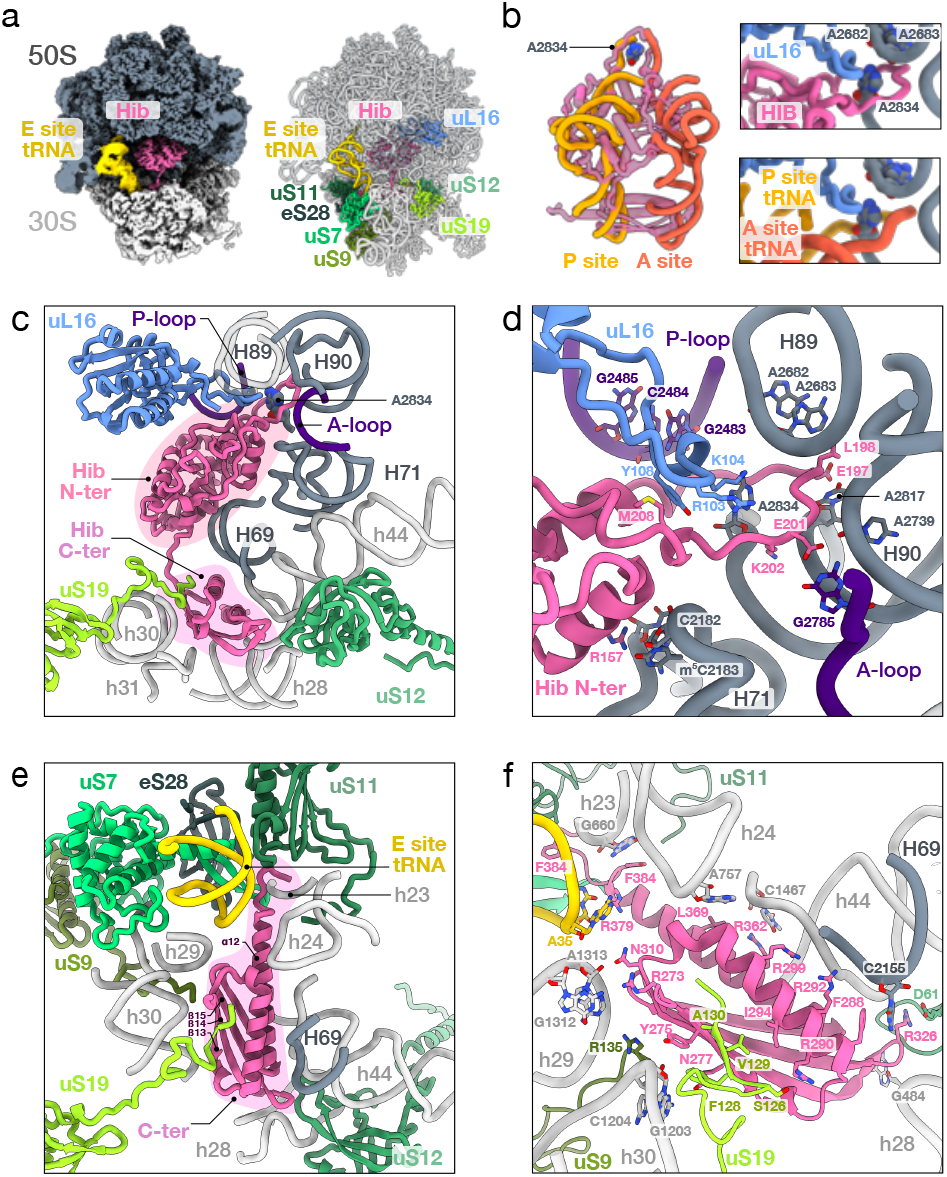
Hib-PTC. a. Hib-PTC 2.1 Å cryo-EM map (left) and structure (right). The cryo-EM map was clipped to show Hib inside of the ribosome. Hib is showed in pink and the E site tRNA in yellow. Ribosomal proteins interacting with Hib are colored and labeled (right). b. Superimposition of Hib onto the structure of a translating human ribosome^53^. Only A and P site tRNAs are shown. Base A2834 universally conserved, located between both A- and P-tRNA CCA ends, is sandwiched by the Hib-loop. uL16 loop is also shown in blue. c. Hib-PTC binding site. Only parts of the ribosome interacting with Hib are represented. d. Close-up of the Hib N-terminal domain inserted in the PTC. Interacting residues are shown in sticks (SupplementaryData 1). e. Close-up of the Hib C-terminal domain binding site. Some of the residues involved in ribosome:Hib contacts are shown in sticks (see also SupplementaryData 1 and Extended Fig. 3. f. View 2 of the Hib C-terminal domain binding site. Some of the residues involved in ribosome:Hib contacts are shown in sticks (see also SupplementaryData1 and Extended Fig. 3.)

The C-terminal end of Hib is at the entrance of the mRNA exit channel. In the Hib-PTC and Hib-intermediate conformations, the α12 helix also interacts with the anticodon of the E site tRNA (Fig. 3e-f). In the A site, Hib C-terminal domain contacts the B2a intersubunit bridge, formed by the interaction between helix-loop 69 (H69) of the 23S rRNA and the upper part of helix h44 of the 16S rRNA. The β14-β15 hairpin loop of Hib interacts with uS12. This loop also contacts the nucleotide h44-A1461 (ec-h44-A1492 *E. coli* numbering) in the decoding site and contributes to stabilize stacking of h44-A1462 onto h44-A1461 and H69-A2154 (ec-H69-A1913 ExtendedData Fig. 3). In addition, this region interacts with G496 (ec-h44-G530). Notably, these four bases are universally conserved and known to be crucial for tRNA substrate selection during decoding^32^. On the other hand, a remarkable interaction involves β13 and the C-terminal tail of uS19, forming a β strand antiparallel to Hib β13 thereby extending the Hib β-sheet (Fig. 3e-f and Supplementary Fig. 4). Finally, contacts are also observed with the C-terminal end of uS9. In total, 270 contacts involving 52 Hib residues were identified (SupplementaryData 1).

We next examined the positions of the Hib N-terminal domain. Cryo-EM analyses revealed three binding sites, almost equally represented in the dataset (Fig. 2). In Hib-PTC conformation (Fig. 3a), a long loop (residues 190-207, hereafter named Hib-loop) is positioned like the CCA ends of A and P tRNAs and surrounds A2834, a universally conserved adenine of the 23S rRNA, sandwiched between the A and P sites on the LSU and known to play a key role in peptide bond formation (Fig. 3a-b and Supplementary Fig. 5). The Hib-loop also engages a large number of interactions with universally conserved nucleotides^33–35^ of the A and P sites at the PTC (Fig. 3d;

SupplementaryData 1). In addition, a long loop of uL16 overhangs Hib-loop and contributes to the stabilization of 23S-A2834. On the other hand, CBS domains 3 and 4 of Hib-PTC interact with the upper part of 23S-H69 while the distal part interacts with the Hib C-terminal domain (Fig. 3c). Overall, positioning of Hib N-terminal domain fully inactivates PTC residues critical for tRNAs binding and catalysis.

In the Hib-intermediate conformation (Fig. 4a), the N-terminal domain rotates by 37° relative to Hib-PTC (Fig. 4c). CBS3 and 4 overlap the elbow and the acceptor helix of the P site tRNA whereas CBS1 and 2 overlap the D and anticodon stems of the E site tRNA (ExtendedData Fig. 4). CBS4 contacts uL5 that makes part of the B1a-B1b/c bridge connecting the 50S to the 30S subunit (Fig. 4a). Only few contacts are observed suggesting that the Hib N-domain can move easily (Fig.4a and SupplementaryData 1).

**Fig. 4:**
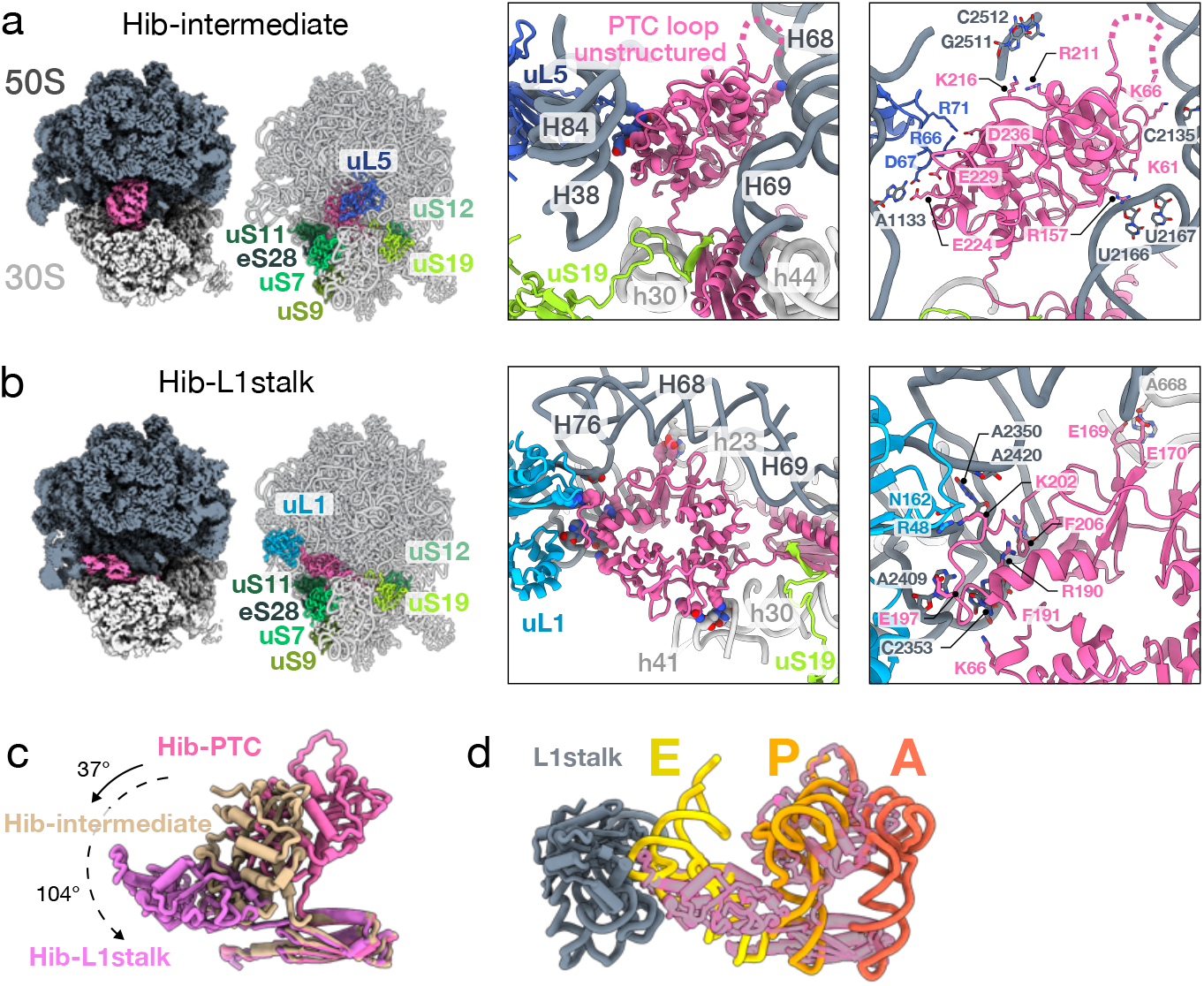
Hib-intermediate and Hib-L1stalk. a. Hib-intermediate 2.0 Å cryo-EM map (left) and structure (right). The cryo-EM map was clipped to show Hib inside of the ribosome. Hib is shown in pink. Ribosomal proteins interacting with Hib are colored and labeled (right). The right-hand views show the interaction of Hib in the B1a-B1b/c bridge region with ul5 and rRNA. b. Hib-L1 stalk 2.1 Å cryo-EM map (left) and structure (right). The right-hand views show the interaction of Hib in the L1-stalk region with ul1 and rRNA. c. Superposition of the three Hib conformations. The rotation angle of the Hib N-domain relative to Hib-PTC is indicated. d. Superposition of the Hib-PTC and Hib-L1stalk conformations on an archaeal translating ribosome with the three tRNAs^54^ (PDB 8HKY). The superposition shows how Hib binding sites overlap those of the tRNAs.

For the third conformation (Hib-L1stalk, Fig. 4b), to improve the quality of the cryo-EM map in the region of the Hib binding site, we performed a local refinement following particles subtraction. The N-domain of Hib rotates by 104° (relative to Hib-PTC) and binds to the L1 stalk through contacts with uL1 and the 23S rRNA (Fig. 4b-d and SupplementaryData 1). The position of the domain corresponds to that of the anticodon helix of an E site tRNA (Fig. 4d and ExtendedData Fig.4). The SSU is rotated as in a post-translocation state.

To conclude, we observe three positions for the N-terminal domain of Hib, while there is no movement of the C-terminal domain. These three positions encompass those of the tRNAs (Fig. 4d and ExtendedData Fig. 4). Thus, Hib acts as a hibernation factor blocking all states of the dynamic translation process, preserving the ribosome from dissociation and degradation.

### Potential partners of the hibernation mechanism in *P. abyssi*

Surprisingly, despite the known propensity of the CBS domains to bind nucleotides, we did not observe any nucleotide bound to the N-terminal domain of Hib in any conformational state. We therefore tested nucleotide-binding by the isolated Hib N-terminal domain using ITC (Supplementary Fig. 6). Hib bound adenylated nucleotides with highest affinity for AMP (Kd=12 ± 2 µM), then ADP (Kd=65 ± 1 µM) and ATP (Kd=113 ± 21 µM). The stoichiometry was consistent with one nucleotide per Hib N-domain (Supplementary Fig. 6). In contrast, no heat signal was obtained when GTP or GMP were used in ITC titrations (data not shown). To assess the importance of nucleotides binding to Hib, we reconstituted ribosomal complexes incubated with 5 mM of either AMP or ATP. Data processing showed that AMP binding did not change classes conformation and distribution (ExtendedData Fig.5). With ATP, the three conformations of Hib were observed, but an additional, weakly populated intermediary state was detected (intermediate2 in ExtendedData Fig. 6). ATP binding was observed in PTC and intermediate states at the interface between CBS3 and 4 (Fig. 5a). Superposition of the free and ATP-bound states showed a slight adjustment of the position of CBS3. The additional intermediate state could reflect higher ribosome:Hib conformational dynamics in the presence of ATP. However, since these experiments were performed at equilibrium with saturating amounts of ATP and in absence of mRNA and tRNA competition, it is difficult to conclude on a potential direct role of ATP in the release of Hib from the ribosome. The role of ATP/AMP binding on CBS module deserves further investigation.

As shown in Fig. 2, SBDS was identified in 4.6 % of the particles as a possible partner of Hib, in the Hib-L1stalk conformation (Fig. 5c). SBDS is a three-domain protein ubiquitous in Eukaryotes and Archaea. In this structure, the N-terminal domain of SBDS occupies the PTC, the second domain interacts with uS12 and the C-terminal domain contacts the P-stalk and the sarcin-ricin loop (Fig. 5c-d). The position of the C-terminal domain overlaps that occupied by the tRNA elbow as observed during the first step of accommodation when the aminoacyl-tRNA:EF1A complex is bound to the ribosome (ExtendedDataFig. 7). Consequently, when Hib binds in the “L1stalk” conformation, the vacant functional sites of the ribosome (PTC, GTPase activating center) can be occupied by SBDS homolog, which thus complements the action of Hib by blocking the ribosome in a completely inactive state that prevents even aa-tRNA:aEF1A initial binding.

**Fig. 5:**
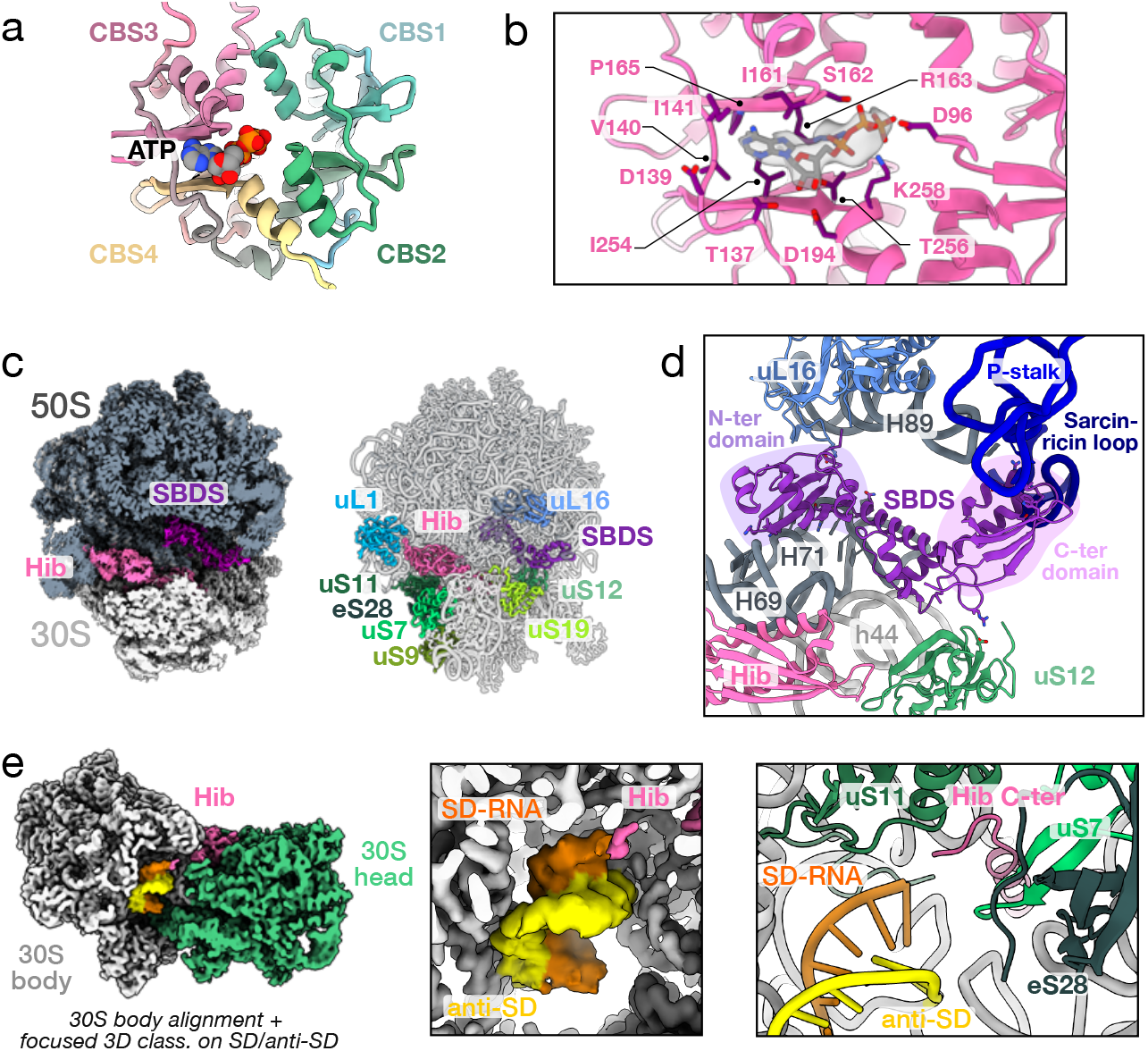
AMP/ATP binding and possible Hib partners. a. Hib bound to ATP in Hib-PTC. b. Residues involved in ATP binding are shown. See also supplementary Fig. 6. c. Hib-L1stalk bound to SBDS 2.4 Å cryo-EM map (left) and structure (right). The cryo-EM map was clipped to show Hib and SBDS inside of the ribosome. Hib is in pink, SBDS is in magenta. Ribosomal proteinsThe N-terminal domain of SBDS interacts with uL16 in the PTC, the intermediate domain contacts uS12 and the C-domain interacts with the P-Stalk and the Sarcin-ricin region. Contacts are detailed in Supplementary Table 1. e. 2.1 Å cryo-EM map from focused classification on SD:antiSD. The SD:antiSD duplex is in orange and yellow. The C-terminal end of Hib is observed in proximity of the SD:antiSD duplex.

**Figure 6:**
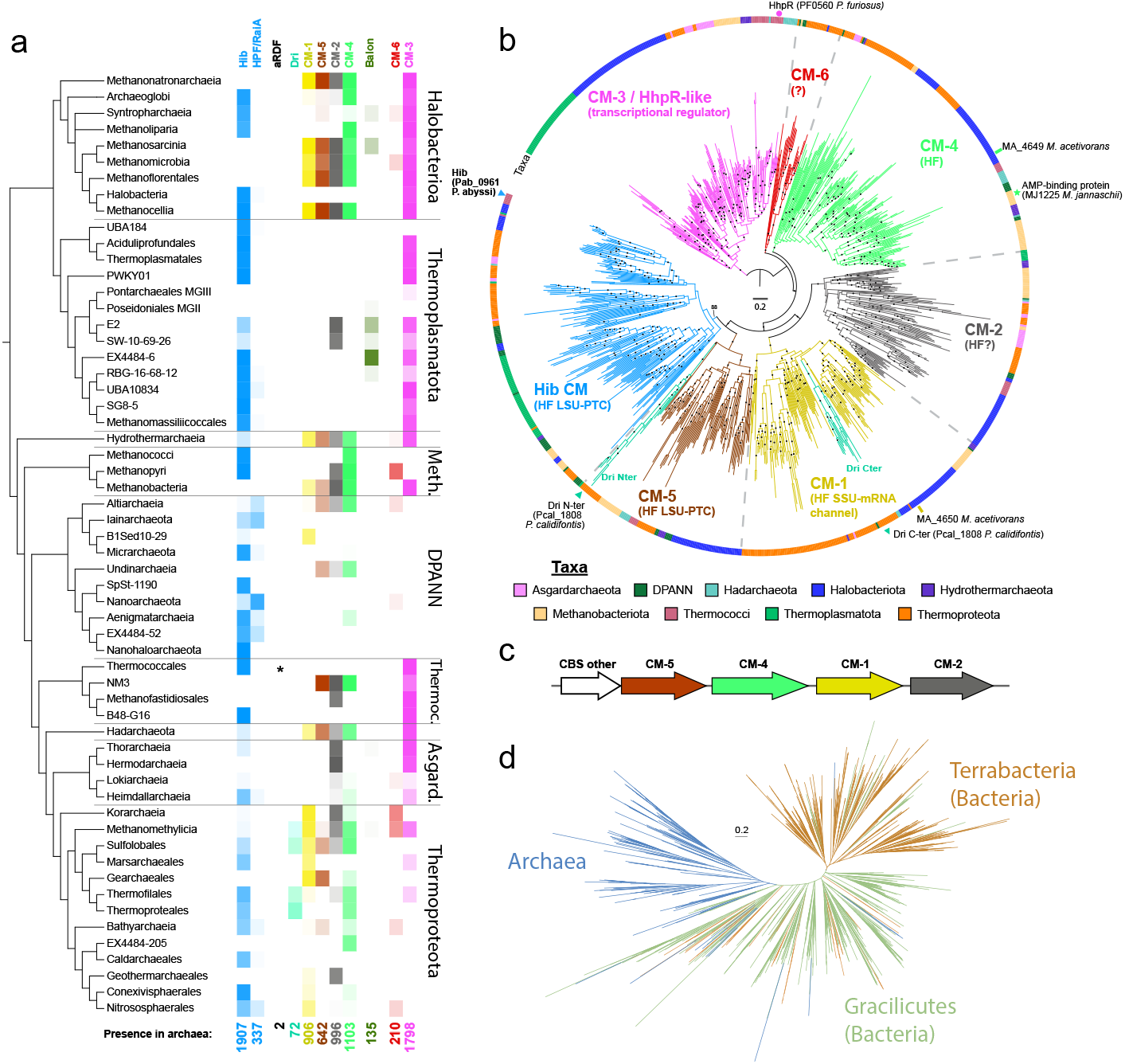
Distribution and phylogenetic relationships between hibernation factors in Archaea. a. Distribution of Hib, Hib-Cter, aRDF, Dri, CBS module (CM) families (CM-1 to CM-6) and potential Balon in Archaea (database of 4026 archaea). The total number of archaea in which these proteins occur is indicated below the heatmap. * indicate the presence of aRDF which otherwise would not be visible with heatmap shading. Meth, *Methanobacteriota*, Asgard, *Asgardarchaeota*. b. Phylogeny of the CM. Branches are coloured according to CM-containing proteins in the panel a. The outer layer indicates the taxonomic origin of the sequences. Position of experimentally characterised proteins are indicated by filled symbol. HF, hibernation factor. SSU-mRNA and LSU-PTC indicates potential interaction sites of the CM members with the ribosome. The tree was rooted on CM-3 which was separated from the other CM by the longest branch and may have different type of interaction (transcription regulator) than the others. Maximum-likelihood tree (LG+R7) based on a trimmed alignment of 230 amino acid positions. Black dotes on the branches indicate ultrafast bootstrap support >90%, with the exception of Hib CM deepest node, value indicated on the tree. c. Example of clustering of CM genes occurring in 10% of the genomes. d. Phylogenie of the HPF/RaiA domain in Archaea and Bacteria.

Finally, during model building, we observed extra density near the anti-SD sequence at the 3’ end of the 16S rRNA. We did focused-classification on the mRNA exit channel (ExtendedData Fig. 2), which revealed the presence of an SD:antiSD helix with 9 paired bases in 17% of the Hib-containing particles (Fig. 5e). This strongly suggests that Hib-mediated ribosome inactivation may involve the binding of an SD sequence forming an SD:antiSD duplex, possibly from a short regulatory RNA or an mRNA degradation product. This SD:antiSD helix may protect the 16S rRNA from its 3’ degradation as reported in bacteria^36^

### Wide distribution of Hib complementing potential alternative hibernation factors in Archaea

Hib was detected in all major lineages of Archaea, corresponding to 47% of all genomes (Fig. 6a and Supplementary Fig. 7). In addition to canonical Hib, we identified proteins composed solely of the HPF/RaiA-like domain. We hereafter refer to these proteins as HPF/RaiA-like, present in 8% of the archaea, mostly DPANN and *Nitrososphaerales* (Fig. 6a and Supplementary Fig. 7). By analogy with Bacteria, these proteins may also contribute to ribosome hibernation.

The absence of Hib from about half of the Archaea, together with the narrow distribution of the recently described Dri and aRDF factors^27,28^ (Fig. 6a), suggested that additional hibernation factors remained to be identified. Both Hib and Dri contain CBS modules (CM), and standalone CM proteins were also identified in dormant ribosome of *Methanosarcina acetivorans* by mass spectrometry^28^. We therefore explored the diversity of CBS modules as potential hibernation factors. Phylogenetic analysis delineated six CBS module families (CM-1 to CM-6) with strong support (Fig. 6b), often arranged in gene clusters (Fig. 6c). Domain placement within this phylogeny indicates that Dri originated from a fusion of CM-1 and CM-5 in *Thermoproteota* (Fig. 6b). Interestingly, CM5 and Hib-CM form a well-supported clade, in agreement with the interactions of both Dri N-terminal and Hib CM with the LSU. Proteins identified in dormant ribosomes of *M. acetivorans* correspond to CM-1 and CM-4. The phyletic distribution of these CM families further provides clues about their involvement in ribosome hibernation. CM-1, CM-2, CM-4 and CM-5 are widespread but rarely co-occur with Hib (Fig. 6a, Supplementary Figs. 7-8). CM-2 and CM-5 in particular, display the strongest patterns of exclusion with Hib. Such exclusion pattern suggests that Archaea generally retain either Hib or CM-1/2/4/5 (fused or standalone) as hibernation factors. In contrast, CM-3 do not display a complementary phyletic pattern with Hib, suggesting that they may not play a role in ribosome hibernation. This is in agreement with the *Pyrococcus yayanosii* HhpR transcription factor, belonging to the CM-3 family ^37^. CM-6 have a narrow distribution, mostly in *Thermoproteota*, and cluster with eukaryotic AMPKγ (Supplementary Fig.9), also suggesting that they are unlikely to be involved in ribosome hibernation.

Finally, we searched for additional homologues of bacterial hibernation factors lacking CBS and/or HPF/RaiA domains. This led to the identification of homologues of the recently described bacterial hibernation factor Balon^14^ in some *Halobacteriota* (*i*.*e*. 30% of the *Methanosarcinia* and 10% of the *Syntropharchaeia*) and several *Thermoplasmatota* lineages (up to 84% in EX4484-6) (Fig. 6a). Their restricted distribution suggests a lineage-specific adaptation. Overall, 84% of the Archaea have Hib, Hib-Cter, Dri, standalone CM-1/2/4/5 and/or the potential Balon.

Among the few lineages lacking identifiable factors, marine planktonic groups are the most notable (*Poseidoniia* (*Thermoplasmatota*) and *Nitrosopumilaceae* (*Thermoprotei*), Fig. 6a), suggesting either a reduced requirement for ribosome hibernation factors in ocean water column or the existence of yet unrecognized hibernation factors.

### Evolution of hibernation factors comprising the HPF/RaiA and CBS modules

The wide distribution of Hib in Archaea reveals that the HPF/RaiA domain contributes to ribosome hibernation beyond Bacteria (Supplementary Fig. 10a-b). Current knowledge of its distribution in Bacteria remains very limited^16^. The discovery of a conserved ribosome hibernation function in Archaea now provides the first opportunity to investigate evolutionary relationships among HPF/RaiA proteins across procaryotes. To this aim, we reconstructed phylogenies of HPF/RaiA-domain proteins in both Archaea and Bacteria and analysed patterns of domain association (ExtendedData Fig. 8).

In Bacteria, the phylogeny of the HPF/RaiA domain (lHPF-NTD, HPF and RaiA proteins, see Supplementary Fig. 1a) reveals a split between the two earliest bacterial branches, *Terrabacteria* and *Gracilicutes*, yet a large distribution across both lineages (Supplementary Fig. 11), consistent with its presence in the Last Bacterial Common Ancestor (LBCA). Mapping of lHPF, HPF/RaiA onto this phylogeny shows that most *Terrabacteria* have lHPF, whereas *Gracilicutes* have both lHPF and HPF/RaiA. Because the CTD of lHPF does not occur as a standalone protein, the current diversity of lHPF is unlikely to reflect multiple independent fusion events. Instead, our data support lHPF as the ancestral bacterial hibernation factor, with the current diversity of HPF/RaiA resulting from multiple losses of lHPF-CTD during Bacterial evolution (ExtendedData Fig. 8).

In Archaea, Hib-CM forms a monophyletic group, closely related to CM-5 (Fig. 6b), consistent with a fusion between an ancestral CM-5 and an HPF/RaiA-like protein. This raises the question of the origin of the “unfused” HPF/RaiA proteins present in current Archaea. HPF/RaiA-like proteins are less common than Hib in Archaea and are generally nested within large Hib clades (Supplementary Fig. 12), supporting multiple emergences through Hib-CM losses. The fusion hypothesis further implies that ribosome binding by standalone CM predates the emergence of Hib. Indeed, CM-5 arose by duplication events that also led to CM-1 and CM-4 (Fig. 6b and ExtendedData Fig. 8), all of which have been detected in dormant ribosomes^28^. Thus, their ancestor may already have played a role in ribosome hibernation.

In the phylogeny of CM-3, all major lineages are represented and monophyletic, suggesting that the duplication separating CM-3 transcriptional regulator from families of hibernation factors occurred before the Last Archaeal Common Ancestor (LACA). This would indicate that ribosome hibernation based on CM proteins lacking the HPF/RaiA domain was present in the LACA.

Finally, we built a combined phylogeny of archaeal (Hib Cter + HPF/RaiA proteins) and bacterial (lHPF NTD + HPF + RaiA) HPF/RaiA domain to infer their phylogenetic relationships. The position of Archaea suggests acquisition of their HPF/RaiA through an early horizontal gene transfer (HGT) from Bacteria (Fig. 6d and ExtendedData Fig. 8). This interpretation should however be taken with caution because of the small size and fast evolution of the domain (<100 aa after trimming). HGT nevertheless remains our preferred hypothesis, as the alternative scenario, where HPF/RaiA-like proteins and standalone CM coexisted in Archaea before fusion with CM-5, is not supported by the current mutual exclusion between Hib, HPF/RaiA-like and CM-5/1/2/4.

## Discussion

Ribosome hibernation factors are central to cellular adaptation under stress. In Archaea, their distribution and physiological roles remain largely unresolved, despite the recent identification of Dri^28^. The discovery and characterization of Hib provide functional insight into the contribution of archaeal ribosome hibernation to stress adaptation. Genetic analysis demonstrates that Hib is critical for efficient recovery from stress in Archaea. In *T. barophilus*, deletion of *hib* led to a pronounced delay in regrowth from stationary phase. In accordance, bacterial ribosome hibernation factors have been consistently shown to be implicated in the ability of cells to resume growth after stress^6^. Mutants lacking HPF or RMF hibernation factors display extended lag phases during recovery from stationary phase^11,12^ or after nutrient starvation^9,38^. Beyond these observations, mechanistic studies suggest that the growth-recovery defect of such mutants can be attributed to a reduction in 70S/100S ribosomes under stress, followed by nucleolytic degradation of ribosomal subunits by the exoribonuclease RNase R^7,8^. This framework is consistent with our finding that the *Δhib* mutant of *T. barophilus* exhibited a marked reduction of 70S ribosomes during stationary phase. By contrast, canonical RNase R homologues are mostly absent from Archaea, and 3’ end ribonucleolytic turnover is primarily mediated by RNA exosome^39^. Together, these data support a conserved role of ribosome hibernation in preserving translation-competent ribosomes under stress. The archaeal Hib factor contributes to ribosome preservation in a manner comparable to bacterial hibernation factors. However, the molecular mechanisms leading to ribosomal degradation might differ between domains and remains to be characterized. The viability assays revealed no detectable difference over long period of stress between the wild type and the *Δhib* strains. In bacteria, the impact of ribosome hibernation on stress period survival is variable. While some studies report reduced viability of hibernation mutants, others find little to no effect, highlighting that the behaviour depends strongly on the stress conditions and genetic background tested (reviewed in^6^). In this instance, it is possible that the conditions assayed may not have been stringent enough to reveal a viability defect, or alternatively that Archaea may possess redundant hibernation factors able to partially compensate for the loss of Hib under prolonged stress. Consistent with its protective role, the expression of *hib* in Thermococcales is controlled by the transcriptional regulator Phr, a master repressor of the archaeal heat shock response^40^. Taken together, its physiological relevance and regulation by Phr place *hib* among archaeal stress-responsive genes, contributing to adaptation across multiple stress conditions.

The results of our cryo-EM study demonstrates that Hib binds the ribosome with three distinct conformations, encompassing the positions of A, P and E site tRNAs during the process of translation. Hence, Hib provides a reversible ribosome inactivation mechanism, being in competition with tRNAs and mRNA bindings when favorable growing conditions return. The three conformations of Hib may illustrate how Hib engages and disengages from the ribosome.

Hib displays a unique modular organization combining a bacterial-type HPF/RaiA core with a CBS module. The C-terminal domain of Hib makes extensive contacts with rRNA and r-proteins in the mRNA channel of the SSU. Comparison with bacterial ribosome structures bound to HPF/RaiA homologs shows that most of the contact points of the hibernation factors with ribosomal RNA are conserved. However, in bacteria, interactions with ribosomal proteins are more limited, and only interactions with uS9 and uS7 have been identified in some cases^14,17,18,29^ (Supplementary Figure 1 and Extended Figure 3). Structural and sequence alignments of the Hib C-terminal domain with bacterial HPF/RaiA proteins show two archaeal-specific insertions absent from most bacterial homologs (Supplementary Fig. 1). One insertion, located between β13 and α11, contributes to the binding site for the C-terminal tail of uS19. In Archaea and Eukaryotes, uS19 protein has an extension of 8-9 residues compared with its bacterial counterpart, and this C-terminal extension interacts with Hib, forming a β-strand antiparallel to Hib β13, thereby extending its β-sheet (Fig. 3d and Supplementary Fig. 1,4). The second insertion, located between β14 and β15, is bound to uS12 and participates in the stabilization of the universally conserved bases crucial for tRNA substrate selection during decoding in the A site^32^. These archaeal-specific insertions illustrate how Hib has evolved to adapt the bacterial-type HPF/RaiA domain to the archaeal/eukaryotic ribosome.

Hib binds to the LSU through its N-terminal CBS module. Orientation of the CBS module defines the three conformations of Hib observed here, in the PTC, bound to the B1a-B1b/c bridge or bound to the L1 stalk. In these three conformations, the CBS module occupies vacant binding sites normally occupied by tRNAs during translation. In the most striking conformation, Hib-PTC, the long Hib-loop completely inactivates the peptidyl-transferase centre. Interestingly, this conformation also involves a specificity of the eukaryotic/archaeal uL16. Although uL16 is universally positioned close to the A-site tRNA binding site, there are notable differences between the eukaryotic/archaeal and bacterial proteins (Supplementary Fig. 13). In particular, eukaryotic and archaeal uL16 have a longer loop inserted into the PTC. The present work shows how the extended loop of uL16 participates in the accommodation of Hib. A similar loop conformation of uL16 may also exist during catalysis, potentially contributing to the positionings of the tRNA CCA-ends and/or substrate accommodation at the PTC^41^. In this view, a comparable uL16 extended loop was modelled in a cryo-EM structure of an archaeal translating ribosome although this point was not discussed in the study^42^.

The importance of CBS modules in hibernation has recently been underscored by the identification of Dri, an archaeal ribosome hibernation factor containing two CBS modules^28^. One CBS module of Dri is observed bound to the PTC. However, the binding of Hib and Dri to PTC differ markedly, with their CBS adopting opposite orientations (ExtendedData Fig. 9). In both cases, however, the long Hib-PTC loop is inserted between CBS3 and CBS4 (ExtendedData Fig. 9). This shows that the versatility of the CBS module binding site on the 50S could be important for the evolution of these hibernation factors. Notably, a long loop located in the same place *i*.*e* between CBS3 and 4, is also present and important for the function in the C-terminal CBS module of Dri and in the recently described stand-alone CBS module protein HhpR, acting as pressure dependent transcriptional regulator in *P. yayanosii* ^37^. Therefore, such insertions could play a recurrent role in the function of these CBS modules.

CBS modules, either standalone or fused with functional domains, are widespread and function as regulatory modules that connect cellular or environmental signals with essential processes. They bind a variety of small ligands, most commonly adenylates but also S-adenosyl methionine, diadenosine polyphosphates or divalent cations, and depending on the ligand may activate or inhibit their protein partner^43^. Here, Hib CBS module preferentially binds AMP, consistent with the general principle that AMP is a sensitive indicator of cellular energy stress ^44^. We observed no clear effect of ATP or AMP on Hib binding to the ribosome. Nevertheless, nucleotide binding could still contribute to Hib engagement or release, or alternatively regulate its interaction with a sequestering partner. In support to this hypothesis, we and others have detected by pulldown approaches coupled to mass spectrometry analyses, interactions between Hib and key actors of DNA recombination and initiation of DNA replication in *Thermococcales* (Hib was referred to as Pab0961 and TK1186 in these studies)^45,46^. In this case, the cytoplasmic concentration of Hib available for its binding to the ribosome would be regulated depending on the ATP/AMP ratio. This possibility remains to be explored.

Together, these observations suggest that the CBS module provides an additional regulatory layer to ribosome hibernation. Yet our structural data also indicate that Hib may act within a broader network of structural or regulatory partners. In the Hib-PTC conformation, the presence of deacylated tRNA in the E site is consistent with the accumulation of uncharged tRNA observed under stress conditions^47^. The E site tRNA may act as a signal to facilitate the recruitment of Hib and to favor the Hib-PTC conformation. In addition to tRNA, SBDS was observed in a subset of Hib-L1stalk particles (Fig. 2c), suggesting an unrecognized function for this factor. In eukaryotes, SBDS is an essential and conserved factor, that controls the final stages of the 60S ribosomal subunit biogenesis, acting in concert with eEFL1 to remove eIF6 from the nascent 60S^48,49^, but such activity has not been observed in Archaea^50^. Here, we found SBDS bound to the ribosome in the context of Hib-L1stalk or associated with an E site tRNA. Its position protects not only the PTC but also the P-stalk and the Sarcin-ricin loop that remain vacant in the absence of aa-tRNA:EF1A. This suggests that SBDS may contribute to ribosome preservation under stress conditions. Interestingly, the conformation observed here corresponds to that of SBDS bound to 60S in Eukaryotes^49^, raising the possibility that SBDS could also interact with the full ribosome in Eukaryotes. Finally, our work suggests that ribosome hibernation in Archaea may also involve the formation of an SD:antiSD duplex that would protect the 3’ end of the 16S rRNA, highly sensitive to RNase-mediated degradation^36^. Thus, ribosome preservation in Thermococcales would be achieved through the protection of all active sites of the ribosome, from the mRNA entry channel to the mRNA exit channel. Altogether, these additional factors highlight that archaeal ribosome hibernation is a multi-layered process, potentially integrating both proteins and RNA-based mechanisms.

Our phylogenetic analyses reveal the evolutionary relationships of HFP/RaiA family proteins across Procaryotes and establish lHPF as the ancestral bacterial hibernation factor. In Archaea, by contrast, the topology and distribution patterns point to an ancestral CM-mediated hibernation mechanism predating Hib, with subsequent domain fusions generating archaeal solutions: the association of an HPF/RaiA-like protein with CM-5 to form Hib, the independent CM-1/CM-5 fusion that produced Dri in *Thermoproteota* and association of unfused CM1 and CM4 in *M. activorans*. Together, these observations highlight the modularity of archaeal hibernation systems, in which CBS modules and HPF/RaiA-like proteins function either as standalone components or in fusion, and illustrate how CBS modules diversified in Archaea to support ribosome hibernation (Hib, Dri) as well as transcriptional regulation (HhpR). More broadly, archaeal CBS proteins may constitute a primordial regulatory toolkit linking cellular energy and environmental signals to essential cellular processes.

These evolutionary insights open perspectives for identifying additional hibernation factors in Archaea. While CM-1, CM-4 and CM-5 have already been detected in dormant ribosomes^28^, the role of CM-2 and CM-6 remains unresolved. Phylogeny and exclusion patterns suggest that CM-2 may also bind the ribosome, whereas the distribution of CM-6 points to a distinct function yet to be explored. In parallel, the detection of potential Balon homologues in Archaea, and its established role as a hibernation factor in Bacteria^14^, support the possibility that Balon-like proteins also contribute to archaeal ribosome hibernation.

Ecological patterns may also highlight niche-dependent specialization of hibernation strategies. Canonical factors are absent from dominant planktonic groups, suggesting that the relatively stable and nutrient-balanced conditions of the ocean water column reduce the selective pressure for such mechanisms. A comparable trend is observed among methanogens: CM-1 and CM-5 (corresponding to the N- and C-terminal moieties of Dri) are specifically absent from gut-associated species (*Methanobrevibacter, Methanosphaera, Methanocorpusculum, Methanomicrobium*) but retained in their closest environmental relative species carrying the CM-5/4/1/2 cluster. This contrast indicates that hibernation strategies involving CM-1 and CM-5, are more important in fluctuating, nutrient-poor environments than in the stable, nutrient-rich conditions of the gut. Accounting for microbial diversity and ecology will be essential for future investigations of Archaeal hibernation strategies.

In conclusion, this study closes a major gap by establishing ribosome hibernation as an adaptive strategy to stress in Archaea, thereby extending this mechanism to all three domains of life. Hib emerges as a key factor in this process, linking archaeal hibernation to the conserved HPF/RaiA domain central to prokaryotic strategies. We also highlight that Archaeal ribosome hibernation has been shaped by CBS modularity and ribosomal plasticity during evolution. Furthermore, this work provides a solid foundation for future work aimed at investigating the role of the potential partners identified, the integration of ribosome hibernation into global stress and energy networks, and the candidate additional ribosome factors in Archaea, yet to be experimentally tested.

## Methods

### Strains, media, and growth conditions

Bacterial and archaeal strains are listed in Supplementary Table 1. *E. coli* strain DH5α was used as the general cloning host. This bacterium was cultivated in Luria-Bertani (LB) broth. Growth of *T. barophilus* was performed in Thermococcales rich medium (TRM), under anaerobic condition at 85°C as previously described^1^. When necessary, media were supplemented with ampicillin (25 μg/mL) for *E. coli*, simvastatin (2.5 μg/mL) and 6MP (100 µM) for *T. barophilus*. Depending on the experiment, elemental or colloidal sulfur (0.1 % or 0.5 g/L final concentration) was added to TRM. Solid media were obtained by adding 16 g/L of agar for *E. coli* and 10 g/L of phytagel for *T. barophilus*. To monitor archaea abundances in liquid media by flow cytometry, an Agilent Advanteon cytometer equipped with a laser with an excitation wave-length of 488 nm was used. Samples (200 μL of culture) were fixed with glutaraldehyde (0.5 % final concentration). After 15 min at 4 °C, the samples were stored at − 80 °C until analysis. The thawed samples were then diluted from 10 to 1000-fold with autoclaved 0.2 μm filtered salted milliQ water solution (NaCl 20 g/L) and stained with the nucleic acid specific dye SYBR®Green I (Invitrogen-Molecular Probes; final concentration: 1) for 15 min at room temperature in the dark. The trigger was set on green fluorescence. Samples were delivered at a rate of 5 μL/min and analyzed during at least 1 min. Salted milliQ water solution and non-inoculated TRM medium were used as negative controls. The data were processed using the NovoExpress tool (v1.4.1).

### Plasmids construction

Primers used are listed in Supplementary Table 2. Deletion of *hib* (TERMP_00697) was performed using plasmid pRD236. This plasmid was constructed by fusion of PCR products obtained using primer pairs 635/636 and 637/638, followed by amplification with primers 635/638. The resulting fusion fragment was inserted, after digestion by *Bgl*II and *Kpn*I, into pUPH previously digested by *Kpn*I and *Bam*HI.

### Transformation methods and strains verification

The transformation of *T. barophilus* (genetic strain Δ*TERMP_00517* will be referred as wild type [WT]^2^) was performed as previously described^3^ using 0.2-2 µg of plasmid. Verification of the deletion was performed by PCR using primer pairs 7/8 and 398/399 to ensure that the non-replicative plasmid used to construct the mutant was not retained in the cell, and with primers 339/340 to confirm the deletion of *hib*.

### Most Probable Number assays

The concentration of cells capable of resuming growth after nutrient stress was determined using the serial dilution culture method, specifically the three-tubes Most Probable Number (MPN) assay^4^. Cultures of 50 mL of both strains were started from precultures and incubated at 85°C until the stationary phase. Cells were then centrifuged for 10 minutes at 8,000 rpm, and washed with carbon-free TRM to remove residual nutrients. After washing, cells were resuspended in 15 mL of TRM lacking carbon sources, and cell concentration was determined by optical cell counting. Subsequently, a 50 mL volume of starvation TRM was inoculated to obtain an initial density of 2,5 x 10^6^ cells/mL. These cultures were then incubated at 85°C, and after 0 h, 5 h, 24 h, 48 h, and 120 h, 6 serial tenfold dilutions in a TRM-rich medium with colloidal sulfur to assess the ability of the WT and mutant strains to resume growth in rich medium after stress exposure. The results are compared with the MPN statistical table^4^ to estimate the concentration of viable cells in the stressed cultures. This method provides an estimate of the number of viable cells with a defined margin of error.

### Sucrose gradient fractionation of whole-cell extracts

*T. barophilus* WT and Δ*hib* cellular extracts were prepared from cells cultivated anaerobically at 85°C under atmospheric pressure (0.1 MPa) in TRM medium^1^, and collected in stationary phase (15 h; 0.5 to 1.108 cells/mL) by filtration (0.45 µm) and centrifugation (26,000 x g for 5 minutes at 4°C). Following a washing step in 1X PBS buffer, 250 mg of cells were re-suspended with 500 mL of SEC100 buffer (20 mM HEPES, pH 7.5; 100 mM NH4Cl; 20 mM Mg acetate; 2.5 mM DTT) containing an EDTA-free protease inhibitor cocktail (cOmplete™, Roche). Whole-cell extracts were prepared by disrupting the cells with one volume of glass beads (>100 µm, Sigma G4649) using two cycles of 2 x 30 seconds at 5,500 rpm at 4°C (Precellys homogenizer). The glass beads were then removed by centrifugation at 5,000 g for 5 minutes. Cell lysates were cleared by centrifugation at 16,000 x g for 10 min at 4°C, followed by filtration through a 0.45 µm Ultrafree-MC HV centrifugal filter (Millipore). The cell extracts were quantified by Bradford protein assay. 1.5 mg of protein was layered onto a linear 10-50 % sucrose gradient in SEC100 buffer. The gradients were then centrifuged at 36,000 rpm in a Beckman SW41Ti rotor for two and a half hours at 4°C. The sucrose gradients were analyzed on a density gradient fractionation system (ISCO UA-6 detector/Brandel Foxy Gradient) with continuous monitoring at 254 nm, allowing the various ribosomal peaks to be resolved.

### Production and purification of *P. abyssi* (His)6-Hib

Hib with an N-terminal (His)6 tag was produced in *E. coli Rosetta*. Cultures (1L) were done at 37°C in 2xYT medium containing ampicillin (50 mg/L) and chloramphenicol (34 mg/L). When OD600nm reached 0.7, expression was induced with 0.5 mM of IPTG and the culture was transferred to 18°C. Cells were lysed by sonication in buffer A (10 mM MOPS-NaOH pH6,7; 500 mM NaCl; 3 mM β-mercaptoethanol) supplemented with PMSF and benzamidine. The clarified lysate was incubated for 5 min at 65°C to precipitate *E. coli* proteins. Hib was then purified by Talon Metal affinity chromatography (Clontech) followed by size exclusion chromatography on a Superdex 200 column (10 mm x 300 mm; Cytiva) equilibrated in buffer A.

### Size-exclusion chromatography of ribosomal complexes

500 pmol of Pa-(His)6-Hib were incubated with 35 pmol of *P. abyssi* ribosomal subunits^5,6^ (30S, 50S or a 1:1 30S:50S mixture) for 45 min at 51°C in 50 µL of ribosome buffer (20 mM Hepes, pH 7.5; 100 mM NH4Cl; 10.5 mM MgAc; 3 mM β-mercaptoethanol). The mixture was then loaded onto an Agilent Bio-SEC5 column (1000Å) equilibrated with ribosome buffer.

### *In vitro* reconstitution of Hib:ribosome complexes

30S and 50S subunits, purified as described in^5,6^, were activated for 5 min at 51°C. 10 pmol of each subunit were incubated for 1 h at room temperature with 50 pmol of (His)6-Hib in a total volume of 12 µL of ribosome buffer for 1h at room temperature. Samples were then diluted to A260 = 4.2 before deposition onto cryo-EM grids. For the Hib-ATP and Hib-AMP datasets, 5 mM of the respective nucleotides and 5 mM MgCl2 were added at the final step of Hib:ribosome incubation.

### Purification of 70S ribosomes from P. abyssi cell extracts

*P. abyssi* cells (DSM 25543)^7^ were grown anaerobically at 90°C and harvested at the end of exponential growth phase (Td = 33 min) as previously described^5^. Cultures were then exposed to aerobic environment and cooled for 1 h in a 4°C cold room before centrifugation. Cell pellets were frozen in liquid nitrogen and stored at -80°C. To purify ribosome cells (0.3 g) were resuspended in 660 µL of ribosome buffer supplemented with RNaseOUT inhibitor (Invitrogen). After addition of an equal volume of 5 µm glass beads (Sigma), cells were vortexed for 5 min. The lysate was centrifuged for 5 min at 5,000 x g and further clarified by two additional centrifugation steps at 19,000 x g (10 min and 5 min) The extract was then fractionated on a 10-40% sucrose gradient centrifuged for 2 h 30 at 36,000 rpm (SW41 rotor, Beckmann). Fractions corresponding to 70S ribosomes were pooled. Sucrose was removed by successive dilution/concentration steps using a Vivaspin 500 PES 30 kDa concentrator, ensuring a residual sucrose concentration below 1% (w/v). Samples were diluted to A260 = 3 for cryo-EM grids preparation.

### Cryo-EM analysis

Samples (3.4 μL) were spotted onto grids (Quantifoil R2/1 Copper 300 mesh with an additional 2 nm continuous carbon layer) at 20 °C and 90% humidity for 10 s. Each sample was vitrified by plunging into liquid ethane at −182 °C, after 1.2 s of blotting using a Leica EM-GP plunger. Conditions were optimized using a Titan Themis cryo-microscope (ThermoFisher) at the CIMEX facility of Ecole Polytechnique. Final datasets were collected on Titan Krios cryo-microscopes (ThermoFisher) available at the ESRF (CM01) or SOLEIL (Polaris) facilities (Supplementary Table 3). Data processing was conducted in Cryosparc 4^8^. The same workflow was used for all datasets described in this study. Briefly, initial 2D templates were generated via blob picking and used to train a Topaz model^9^. Topaz picks were then used to generate three initial 3D models, which were refined through heterogeneous refinement. Further 3D classifications, global or focused, were performed before final homogeneous refinements. Subsequent global and focused 3D classifications were carried out prior to final non-uniform refinements. Flexible regions were further refined using 3D Flex^10^ to improve local map quality. A detailed description of all processing steps is provided in Extended Data Figs. 1,2,5 and 6. Particle orientation diagnostic and local resolution cryo-EM maps are shown in Supplementary Fig. 14.

Models were built in Coot^11^ and refined in Phenix^12^. 3D structures of the 30S from *P. abyssi*^5,6,13^ and that of the 50S from *T. kodakarensis* and *P. furiosus*^14,15^ were used as starting models. Examples showing the quality of the cryo-EM map for rRNA modified nucleotides (Supplementary Table 4), magnesium ions, Hib environment are given in Supplementary Fig. 15 and 16. Water molecules were only added in the Hib-PTC reference structure (2.1 Å, PDB 9SRE). rRNA secondary structure diagrams are given in Supplementary Fig. 17 and 18. They have been updated according to the 3D structures. All structural figures were drawn using ChimeraX^16^.

### Isothermal Titration Calorimetry

Before ITC measurements, Hib N-terminal domain was dialyzed against buffer B (10 mM MOPS-NaOH pH6,7; 50 mM NaCl; 3 mM β-mercaptoethanol) containing 2 mM MgCl2. Titrations curves were obtained using a MicroCal200 apparatus (Malvern). The protein (202 µL) was titrated through 20 injections (4 µL) of ATP (2.48 mM) or ADP (2.45 mM) or AMP (0.76 mM) solutions in buffer B containing 3 mM MgCl2. Concentrations of proteins were 194 µM (ATP and ADP titrations) or 100 µM (AMP titration). Typical titration experiments are shown in Supplementary Fig. 6. Dissociation constants and stoichiometries were deduced from least-square fits of measured heat values to standard binding isotherms using Origin (OriginLab). Reported results are mean ± s.d. from three (ATP or AMP) or two (ADP) experiments. Titration experiments with GTP or GMP (1 mM in the syringe) did not cause any measurable heat change.

### Genome Databases

Two genome databases covering the diversity of archaea were assembled, a minimal database with 388 genomes (Arc388) and a larger one with 4026 genomes (Arc4026). Arc388 was used for phylogenetic analyses while the Arc4026 was used for distribution analyses. All archaeal genomes available in NCBI (2024/01/16) were collected, as well as genomes from other published datasets, including Earth, ocean, soil, glacier and termite MAG catalogues^17–21^. To build the Arc4026 database, these genomes were dereplicated at 92% ANI using dRep^22^, and only genomes with more than 80% completeness and less than 5% contamination according to CheckM were retained^23^. The Arc388 database is a manually curated sub-selection of genomes covering main archaeal lineages. For Bacteria we used a previously published database of 1067 genomes covering all major lineages^24^.

### Phylogeny and distribution of Hib, potential alternative hibernation factors

All sequences were aligned with MAFFT L-INS-i^25^ and trimmed with BMGE-1.12^26^ using the BLOSUM30 substitution matrix. All maximum-likelihood trees were built with IQ-TREE 2.0.6 using the TESTNEW option to define the best model^27^. Tree visualization, colouring, and protein presence/absence mapping were performed with ITOL^28^.

Pa-Hib was used as a seed to search for the closest sequences in Arc4026. From the C-terminal region (corresponding to the RaiA domain of Hib) of these sequences we built an HMM domain using HMMBUILD (hmmer v3.3.2 [http://hmmer.org/]). Additional Hib sequences were identified in Arc4026 using the Hib-Cter HMM profile (HMMSEARCH) and incorporated for iterative refinements of the profile. Specificity of the hits was assessed by identifying additional conserved domains, including the presence of the CBS module, and by examining Alphafold2-predicted protein structures available in Uniprot (https://www.uniprot.org/) for the most divergent sequences. Shorter sequences lacking the CM module but displaying the HPF/RaiA fold were retained.

The Hib alignment was also truncated at the C-terminal region to generate a Hib-Nter HMM profile corresponding to the CBS module. Using this profile, additional proteins containing a CBS module were identified in the Arc388 database. All proteins with an E-value lower than 10^-28^ were retained. Most proteins with a higher E-value are composed of a single tandem of CBS instead of two and can be associated with diverse alternative conserved domains. Selected hits included the Dri hibernation factor, formed by a fusion of two CBS modules. Dri sequences were split and annotated as Dri Nter and Dri Cter. After alignment trimming, a maximum-likelihood tree was built and used to delineate paralogous protein families corresponding to CBS modules (called CM-1 to CM-6), in addition to the family formed by the CBS module region of Hib. The description of HhpR (MJ0729), belonging to CM-3 family was published recently^29^ and its genome was not part of the Arc388 database. The sequence of this protein was thus not retrieved from our analysis but directly added to the CM alignment.

HMM profiles were built for each of these families and used to determine their distribution in the Arc4026 and Bac1067 databases. Alternative hibernation factors known to occur in bacteria (Etta, SRA, ElaB/YqjD, RMF, RsfS and Balon) and Eukaryotes (Stm1, IFRD2, Lso2, MDF1, MDF2 and Dapl1) were searched using available COG or pfam HMM profiles or by BLASTp searches using the originally characterized proteins. For Balon, an HMM profile was generated using proteins identified in Bacteria^30^. With the exception of Balon, few or no significant hits were obtained. Potential archaeal Balon proteins were distinguished from homologous aeRF1 and pelota proteins by identifying sequences lacking the NIKS and GGQ motifs of aeRF1 and forming a distinct monophyletic lineage in a ML phylogeny. All genomes encoding a putative archaeal Balon protein also have aeRF1, and sometime a pelota protein. Identified hibernation factors were mapped on a tree containing all archaea from the Arc4026 database. Markers used to build this tree correspond to the Phylosift dataset^31^ and five additional markers as described in^32^.

Sequences containing the HPF/RaiA domain and the lHPF CTD were searched in the Bac1067 database using the COG1544 and PF16321 HMM profiles, respectively. To identify conserved domains fused to HPF/RaiA we used CD-search^33^ on proteins longer than 150 aa lacking the lHPF CTD. Phylogenies of the bacterial lHPF and HPF/RaiA without addition domain were built with and without the addition of the Hib-Cter domain and HPF/RaiA-like proteins from Archaea.

### Data availability

The atomic models and cryo-EM maps have been deposited to the Protein Data Bank. The PDB and EMDB codes are as follows, 70S-Hib-PTC: 9SRE, EMDB 55139; 70S-Hib-PTC-wo-E-tRNA: 9SRC, EMDB-55137;70S-Hib-intermediate : 9SRD, EMDB-55138; 70S-Hib-L1stalk : XXX; Hib-CBSmodule-L1stalk : 9SRF, EMDB-55140;70S-Hib-L1stalk-SBDS : 9SRB, EMDB-55136;70S-Hib-SD-antiSD : 9SRA, EMDB-55135; Hib-ATP: 9SR9, EMDB-55134.

## Acknowledgements

This work was supported by grants from the Centre National de la Recherche Scientifique and Ecole polytechnique to Unité Mixte de Recherche n°7654 and by a grant from the Agence Nationale de la Recherche (ANR-25-CE12-4161). Cryo-EM data benefited from access to CM01 beamline (ESRF, Grenoble), the Polaris Beamline (SOLEIL synchrotron, Saint Aubin, France) and the Interdisciplinary Center for Electron Microscopy of Ecole Polytechnique (CIMEX). We thank the staff at these facilities, and in particular Romain Linarès (ESRF), Eric Larquet, Pierre Legrand (SOLEIL) and Kassiogé Dembélé (CIMEX). We thank David Mignon for skillful computer assistance.

## Author Contributions

DF, ES, YM, CM, GaB, GuB, MB conceived and designed the study. CM, GaB, CD performed the biochemical experiments and complexes preparation. CM, GaB, YM, ES did cryo-EM analysis, model building and data analysis. MK, RD, KF, CD, SL, LM performed thermococcales cultivation and genetic experiments. MK, BCO, MB, RC performed ribosome sedimentation analyses. GuB performed taxonomic and phylogenomic analyses. DF, ES, YM, GuB, MB drafted the manuscript with input from all authors. All authors reviewed and edited the manuscript.

---

**ExtendedData Fig.1:**
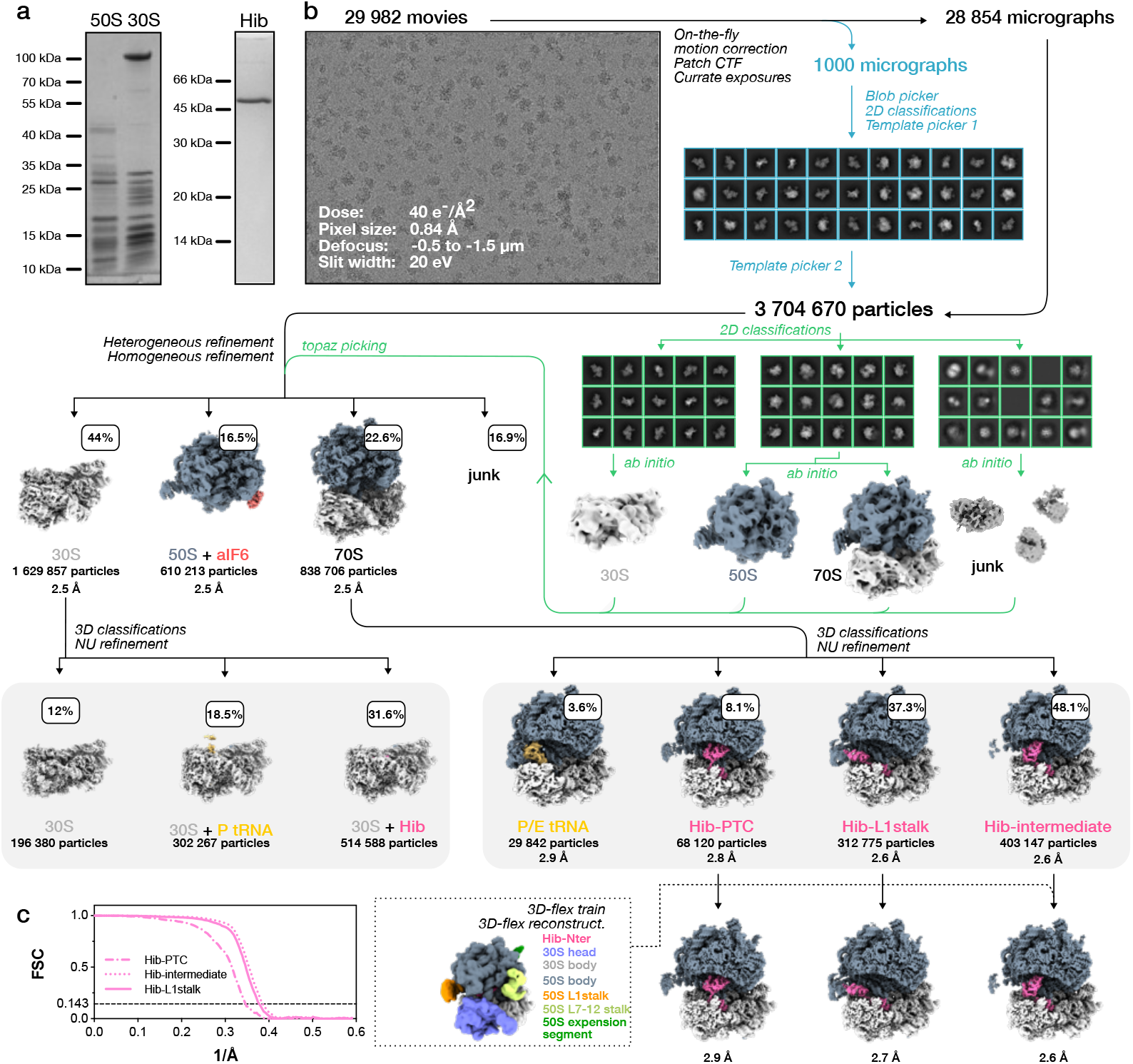
Cryo-EM structure determination of in vitro reconstitued hibernating ribosomes. a. SDS-PAGE analysis of purified *P. abyssi* ribosomal subunits and recombinant Hib. b. Cryo-EM workflow. Image processing was carried out in cryoSPARC v4.1. After motion correction and CTF estimation, micrographs were screened based on CTF fit (<5 Å) and ice thickness. From an initial subset of 1,000 images, particles were picked using the blob picker and classified in 2D to generate templates for large-scale picking. Particles recovered from template-based picking underwent iterative 2D classification, ab initio reconstruction, and heterogeneous refinement to eliminate contaminants. The cleaned particle set was then used to retrain Topaz for particle picking across the full dataset, yielding 3,704,670 particles. Extracted particles (box size 480 pixels) were classified in 3D against ribosome and junk references, and the resulting 70S ribosome particles were subjected to focused classification on the tRNA-binding sites. Hib-bound classes were further refined using non-uniform refinement followed by 3D Flex. Note that free 50S are bound to aIF6, excluding their association with the 30S. The 50S:aIF6 particles correspond to a final stage of 50S biogenesis that co-purify with the mature 50S. c. Resolution of final reconstructions was determined by gold-standard FSC at the 0.143 criterion.

**ExtendedData Fig.2:**
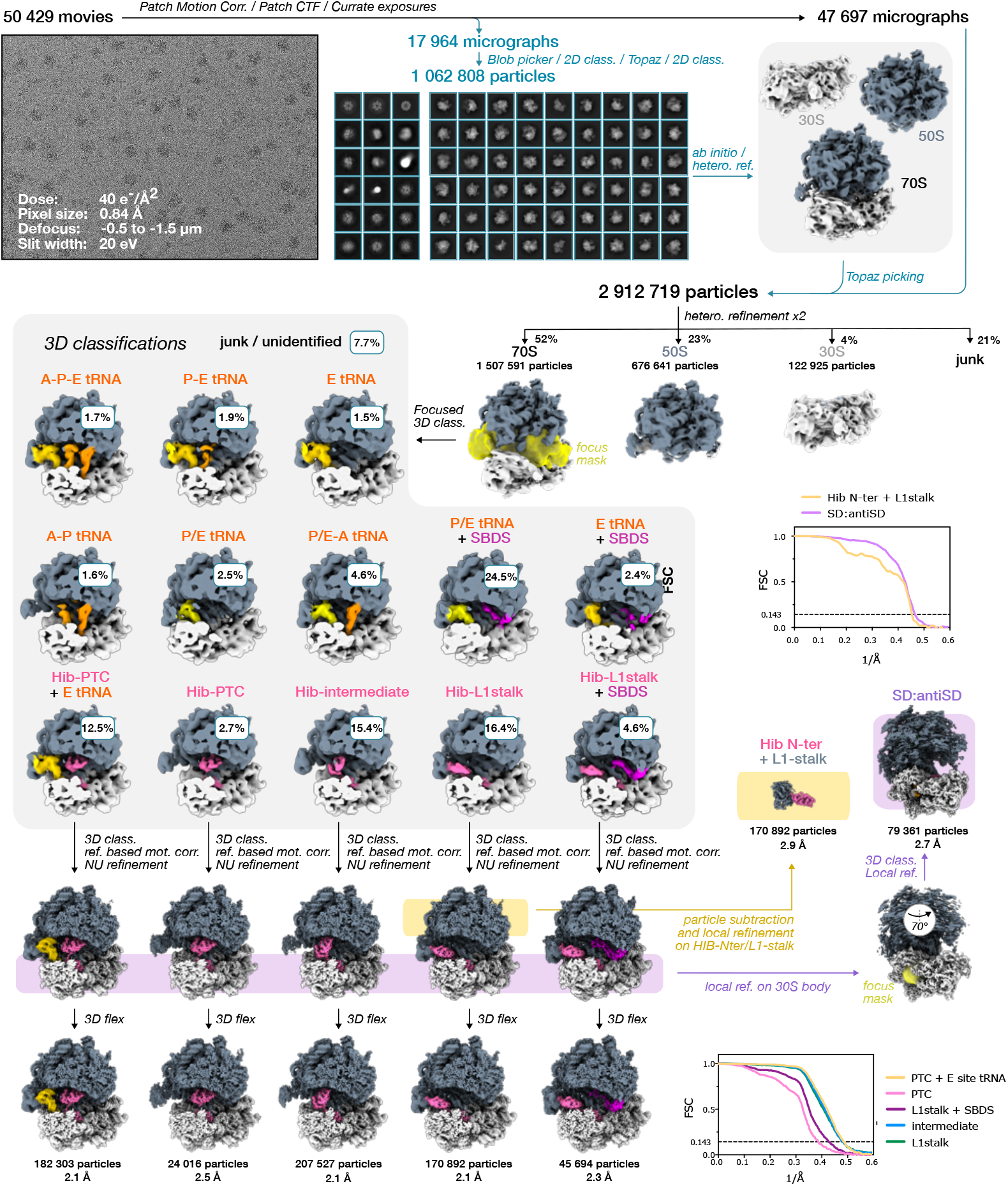
Cryo-EM structure determination of ribosomes from cell extracts. Data processing was performed in CryoSPARC v4.1. Following motion correction and CTF estimation, exposures were filtered based on CTF fit (<5 Å) and ice thickness. An initial subset of 17,964 images was processed by blob-based particle picking and 2D classification to retain ribosome classes, which were used to train Topaz. Particles repicked with the trained model underwent iterative 2D classification, ab initio reconstruction, and heterogeneous refinement to remove contaminants, and the resulting clean subset was used to retrain Topaz for large-scale picking. Using this model, particle picking across the entire dataset yielded 2,912,719 particles (box size of 480 pixels) which were classified in 3D against ribosome and junk references. The resulting 70S ribosomes were subjected to focused classification on the tRNA-binding sites, and Hib-bound classes were refined with reference-based motion correction to generate final maps using 3D Flex, with the masks shown in ExtendedData Fig1. To enhance resolution of the Hib N-terminal/L1 stalk complex, particle subtraction with a mask excluding the rest of the ribosome mask was followed by local refinement centered at the L1 stalk. In parallel, Hib-bound particles were aligned on the 30S body, classified with a mask over the SD:anti-SD region, and the resulting particles were refined with a 30S body mask.

**ExtendedData Fig.3:**
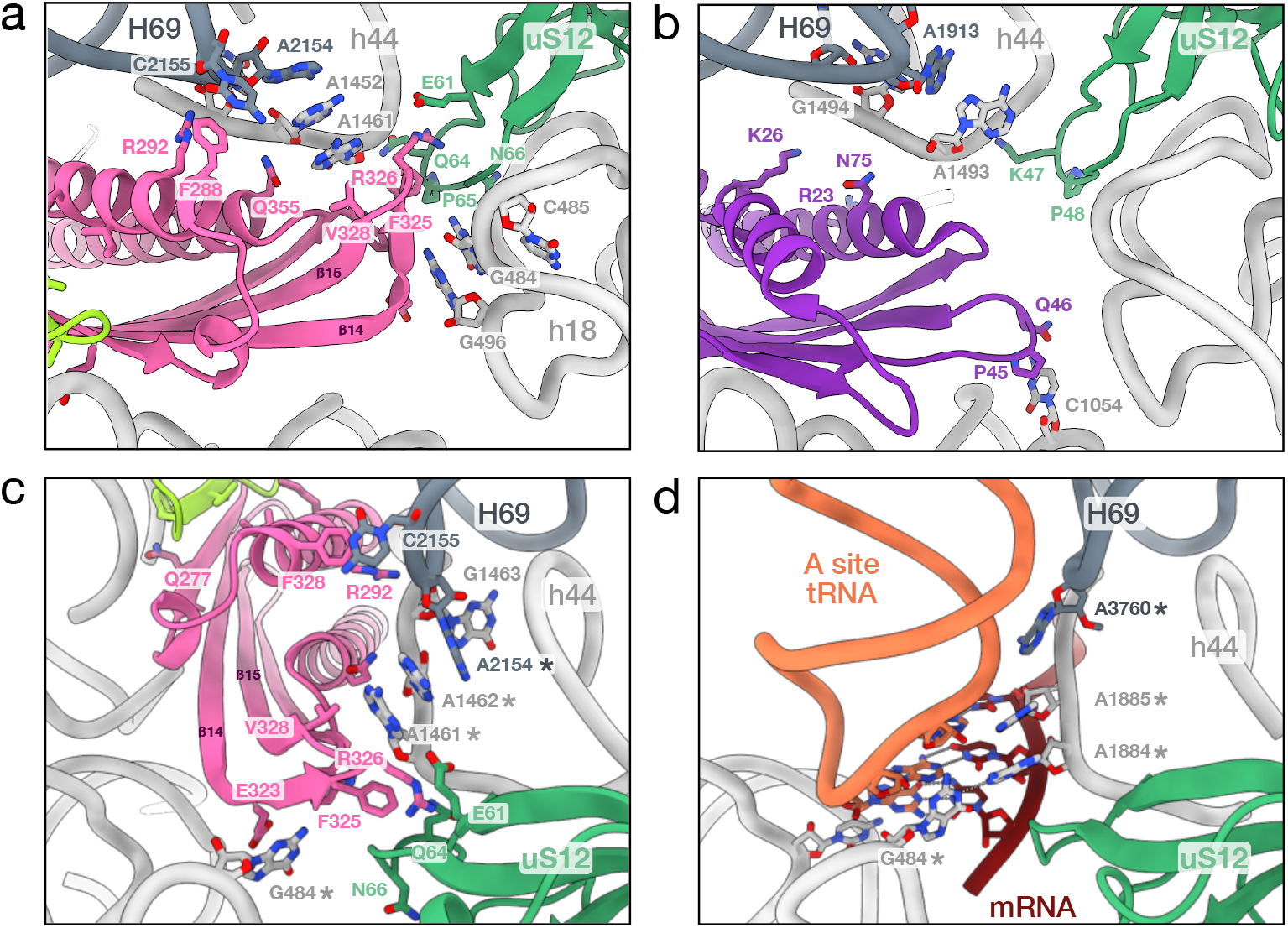
Interaction of the C-terminal domain of Hib with uS12 and the B2a bridge at the A site. a. Close-up of the contacts between uS12 (in green), the B2a bridge (H69:h44) and the C-terminal domain of Hib. Only the residues involved in contacts are shown in sticks. Insertion 325-330 is specific of archaeal HPF/RaiA domains. The contacts were calculated with ChimeraX. b. Structure of *E. coli* RaiA (PDB 4V8I, {Polikanov, 2012 #4784}) bound to the ribosome shown in the same orientation as in view a. Residues involved in contacts between RaiA and the ribosome are shown in sticks. c. Identical to a. but zoomed in on A site. Bases h44-A1461, h44-A1462, h18-G484 and H69-A2154 (corresponding to A1493, A1492, G530 and A1913 *E. coli* numbering) are indicated by a star. d. Same region but in the structure of human ribosome with A site tRNA and mRNA (PDB 8G61, {Holm, 2023 #4638}). The two structures were aligned by superimposing the 16s rRNA (rmsd 1.1 Å for 747 atom pairs). The view shows the codon:anticodon interaction and the position of the four bases universally involved in the decoding step. Comparing views c. and d. shows how Hib interacts with bases important for the decoding mechanism.

**ExtendedData Fig.4:**
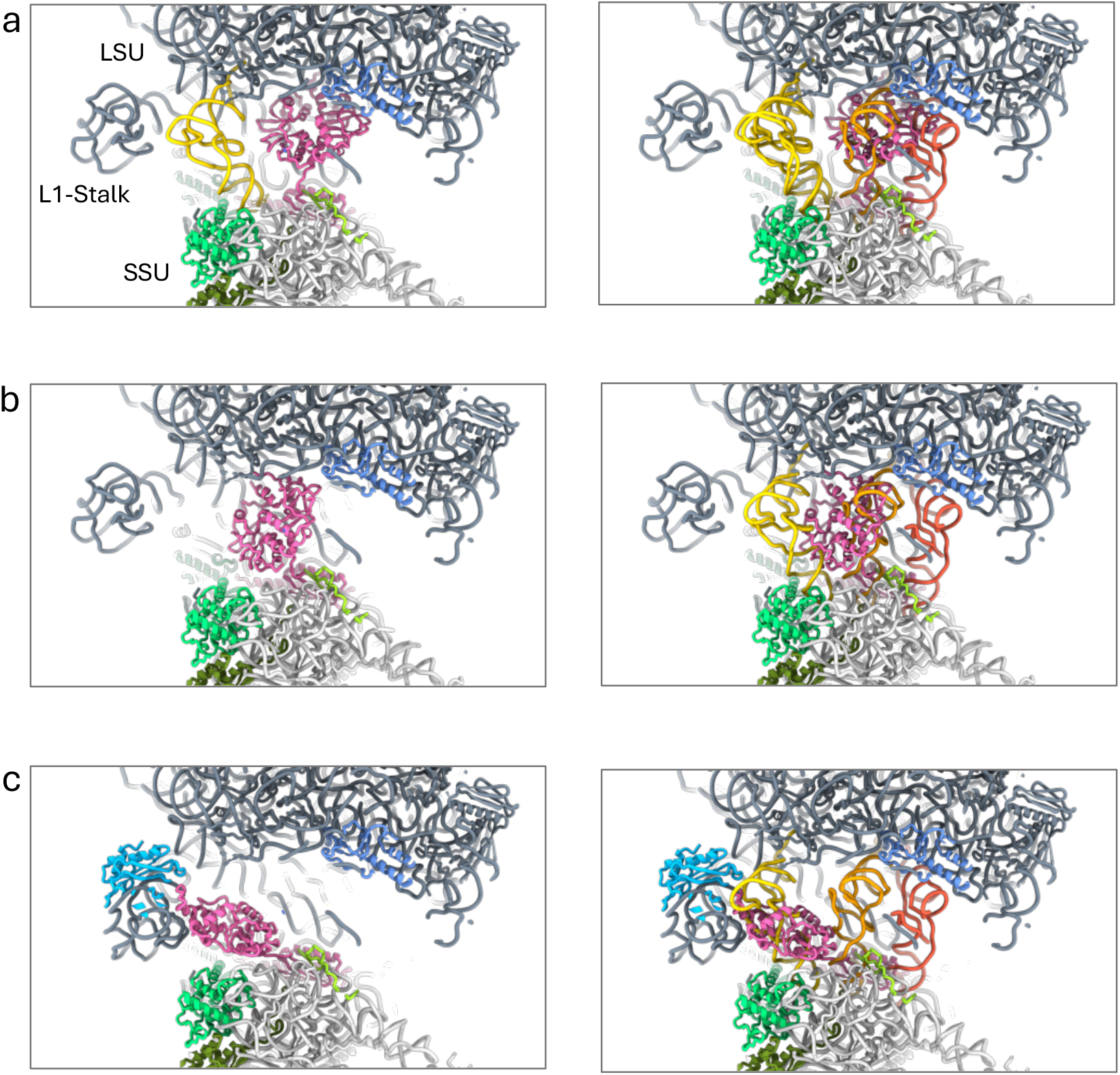
comparison of Hib binding sites with tRNA binding sites. The structures Hib-PTC (a), Hib-intermediate (b) and Hib-L1stalk (c), left panels, were superimposed to the structure of *S. acidocaldarius* ribosome bound to three tRNAs, right panels (PDB 8HKY, {Wang, 2023 #4620}). Color code is as follows: E site tRNA gold, P site tRNA orange, A site tRNA tomato. Hib is in pink. uL16 is in cornflower blue, uL1 is in light blue, uS19 is in light green and uS7 in green.

**ExtendedData Fig.5:**
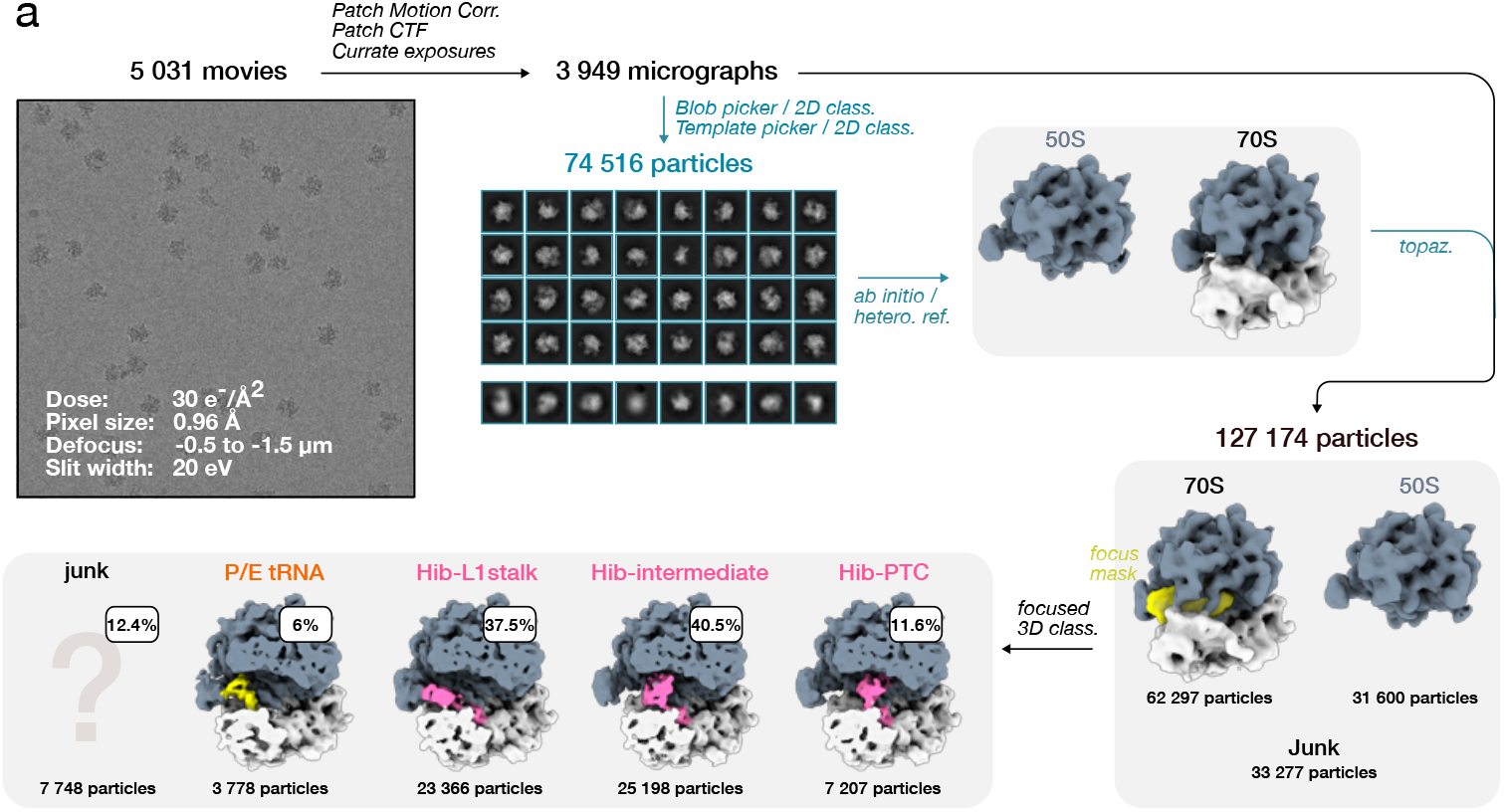
Cryo-EM structure determination of in vitro reconstitued hibernating ribosomes in the presence of AMP. Image processing was performed in cryoSPARC v4.1. Following motion correction and CTF estimation, micrographs were filtered based on CTF fit (<5 Å) and ice thickness. Particles were initially picked with the blob picker and classified in 2D to generate templates for subsequent particle picking. Particles obtained from template-based picking were subjected to iterative 2D classification, ab initio reconstruction, and heterogeneous refinement to remove contaminants. The resulting clean subset was then used to train Topaz for particle picking across the dataset, yielding 74,516 particles. Extracted particles (box size 480 pixels) were classified in 3D against ribosome and junk references, and the resulting 70S ribosomes were further analyzed by 3D classification focused on tRNA-binding sites. AMP binding was observed in the cryo-EM maps but due to the low number of particles no attempt was made to build a full model.

**ExtendedData Fig.6:**
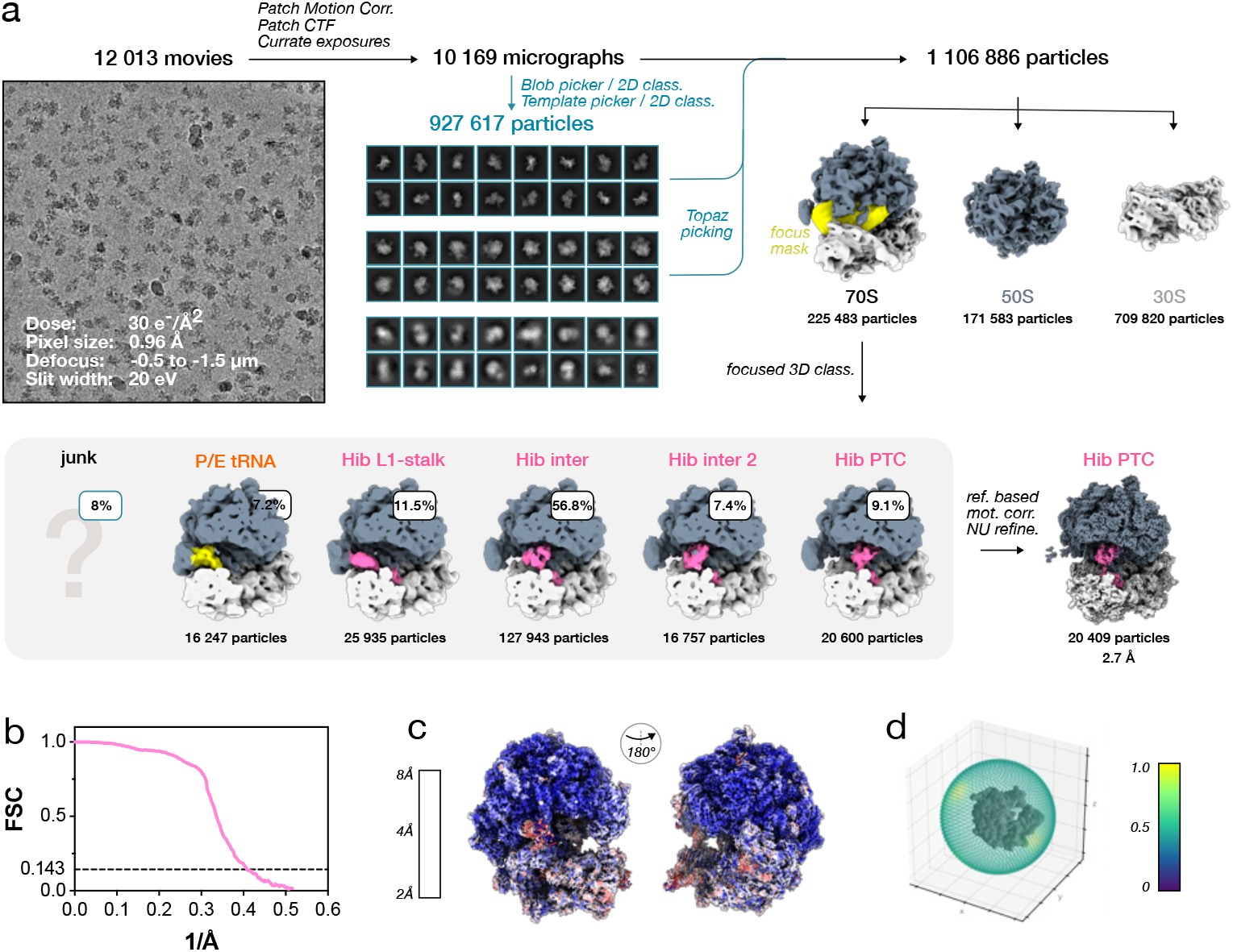
Cryo-EM structure determination of in vitro reconstitued hibernating ribosomes in the presence of ATP. a. Cryo-EM workflow. Data processing was performed in cryoSPARC v4.1. After motion correction and CTF estimation, exposures were filtered based on CTF fit (<5 Å) and ice thickness. Particles were initially picked with the blob picker and classified in 2D to generate templates for particle picking. Newly picked particles were classified in 2D, and ribosome particles were used to train Topaz. Particles repicked with the trained model were extracted in 420-pixel boxes and classified in 3D against ribosome and junk references. The resulting 70S ribosomes were subjected to focused classification on the tRNA-binding sites, and the Hib-PTC conformation was further refined following reference-based motion correction. b. Resolution of the final reconstructions was determined by gold-standard FSC at the 0.143 criterion. c. Local resolution of the final reconstruction. d. 3D scatter plot.

**ExtendedData Fig.7:**
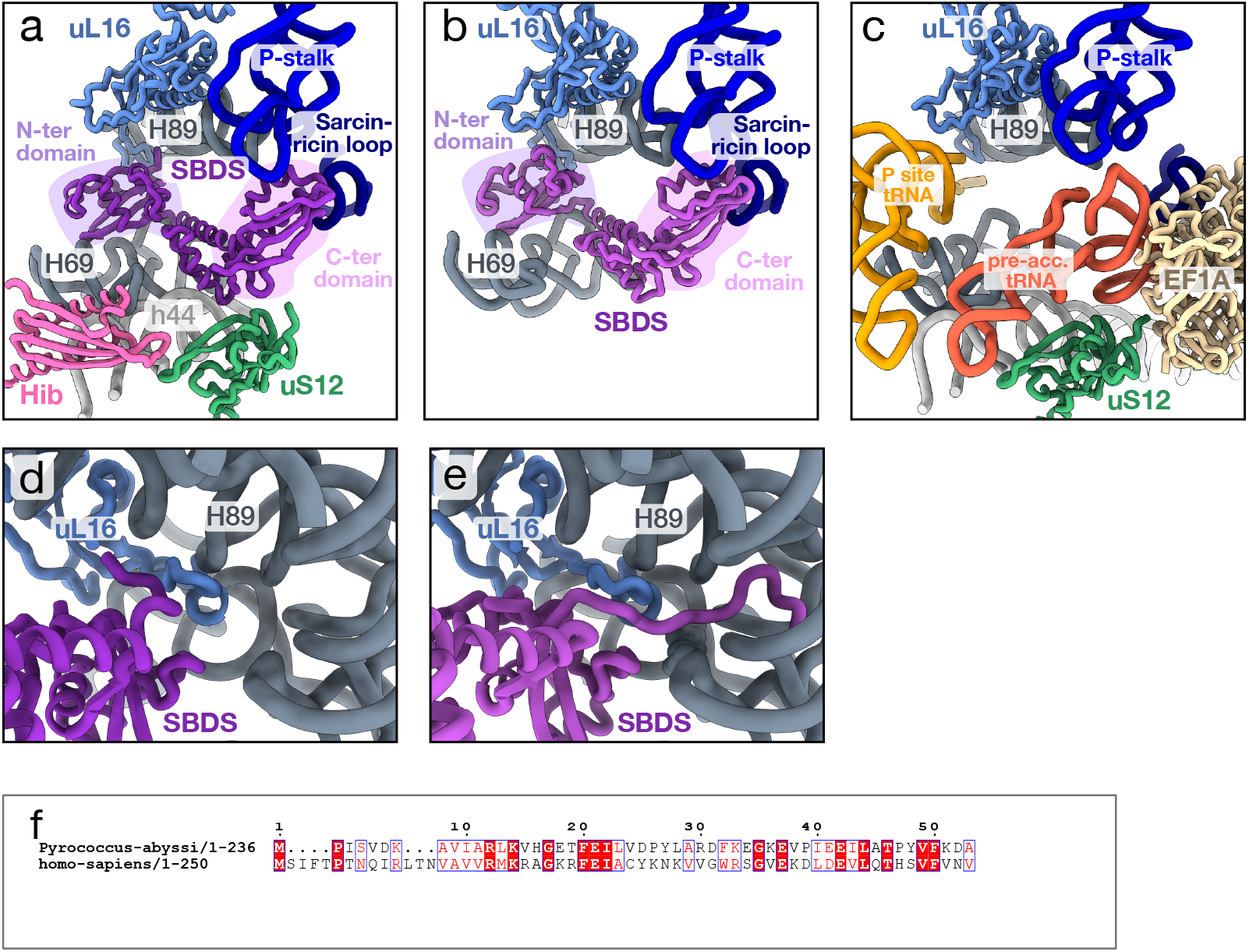
Binding site of SBDS to the ribosome. a. Binding of Pa-SBDS to Hib-L1stalk. rRNA regions and r-proteins interacting with SBDS are shown and labeled. b. Binding site of Human SBDS to *Dictyostelium discoïdeum* 50S (PDB 6QKL, {Weis, 2015 #4792}). Comparison of views a. and b. shows that the binding sites of SBDS to the ribosome or to the LSU are very similar. b. Same orientation as in view a. but for human translating ribosome with a pre-accommodated aa-tRNA:EF1A complex (PDB 8G5Z, {Holm, 2023 #4638}). The view shows that the C-terminal domain of SBDS occupies the same position as that of the elbow of a pre-accommodated aa-tRNA. c. Same as a. but zoomed on the PTC. The view shows the orientation of the N-terminal end of Pa-SBDS. d. Same as b. but zoomed on the PTC. The view shows the orientation of the N-terminal end of Human SBDS in the peptide exit tunnel. e. Sequence alignement of Pa-SBDS and human SBDS. The archaeal versions of the protein are systematically 6 to 7 residues shorter in their N-terminal end as compared to eukaryotic SBDS versions. This may explain why the N-terminal end of Pa-SBDS is not observed in the peptide exit tunnel (compare views d. and e.). For clarity, the alignement of Pa-SBDS and Hs-SBDS sequences is only shown for the first 50 residues.

**ExtendedData Fig.8:**
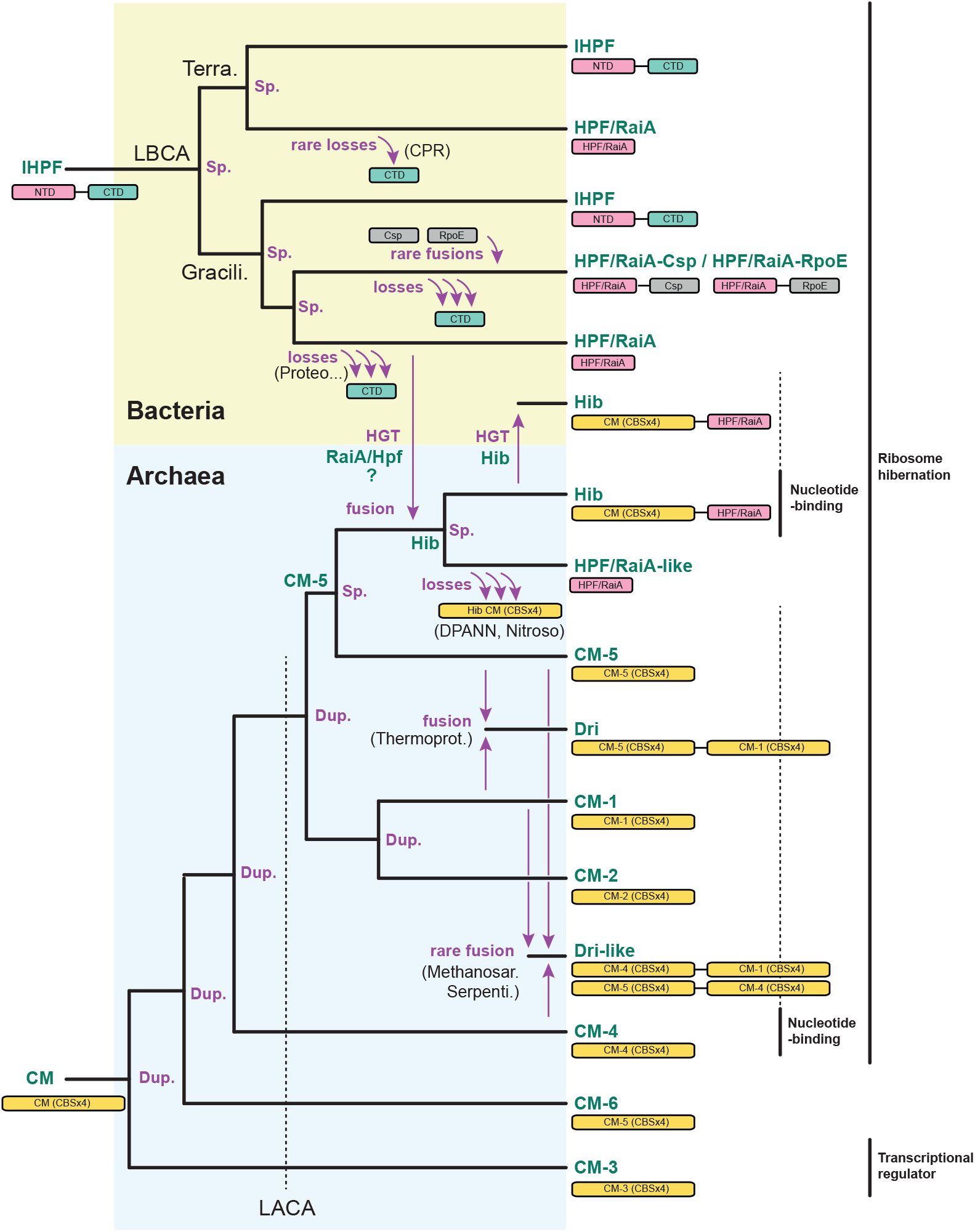
Evolutionary scenario of proteins containing the RaiA/HPF domain and CBS modules (CM) in Archaea and Bacteria. Sp., speciation; Dup., duplication; Proteo., Proteobacteria; Nitroso., Nitrososphaerales; Thermoprot., Thermoproteota; Methanosar., Methanosarcinaceae; Serpenti., Serpentinarchaeaceae. See text for description.

**ExtendedData Fig.9:**
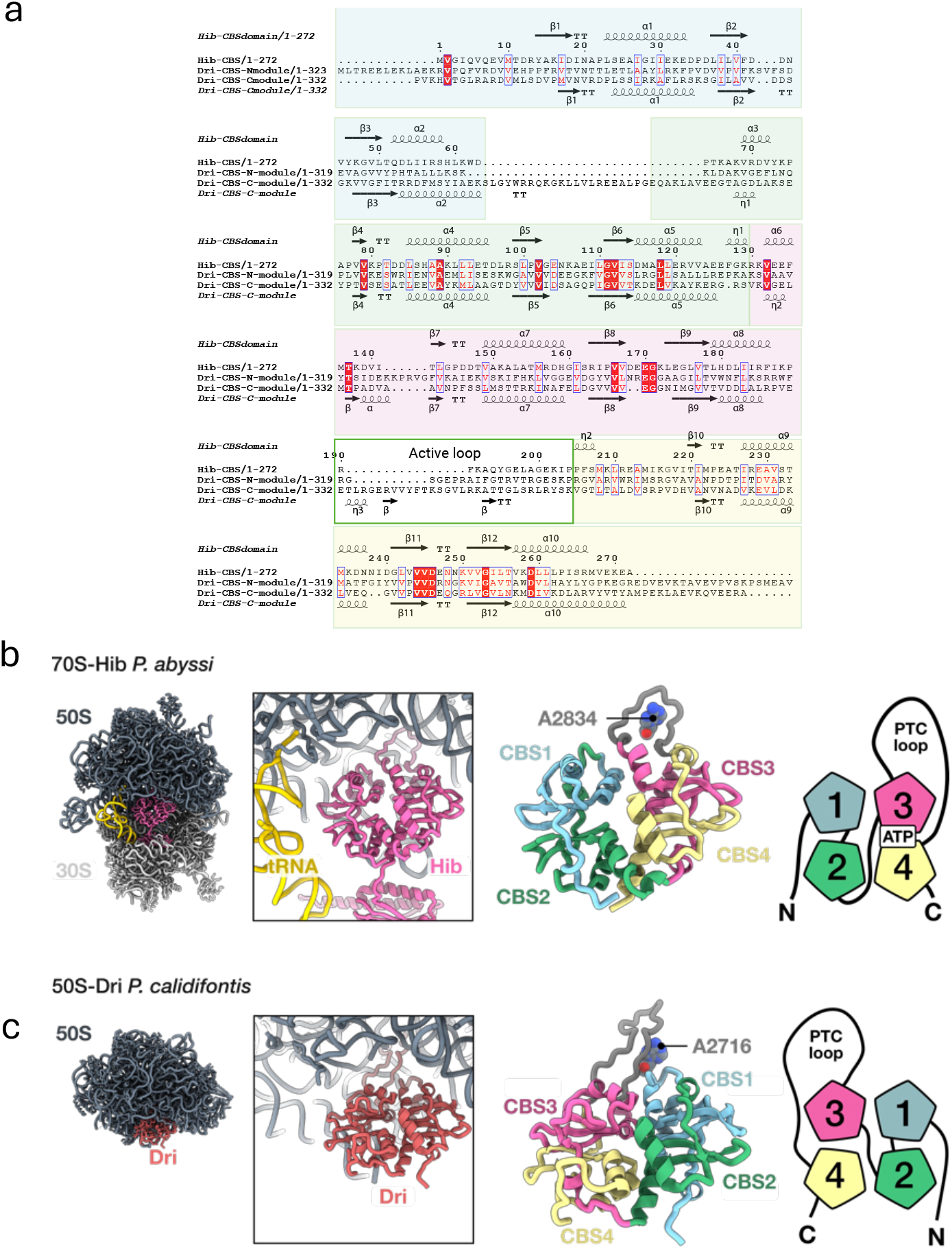
Comparison of Hib and Dri CBS modules. a. Structural alignement of the CBS modules from Hib and Dri (N and C). The CBS domains are highlighted as follows; CBS1 blue; CBS 2 green, CBS 3 pink, CBS 4 yellow. b. View of Hib-PTC. On the right side, a diagram shows the arrangement of the CBS domains. c. View of Dri N-CBS module bound to the 50S (PDB 9E6Q, {Nissley, 2025 #4798}). A diagram shows the arrangement of the CBS domains. In views b and c, the 50S subunits have been superimposed. Comparison of views b. and c. shows that Hib and Dri (N-module) orientations differ by 180°.

**Supplementary Fig. 1.**
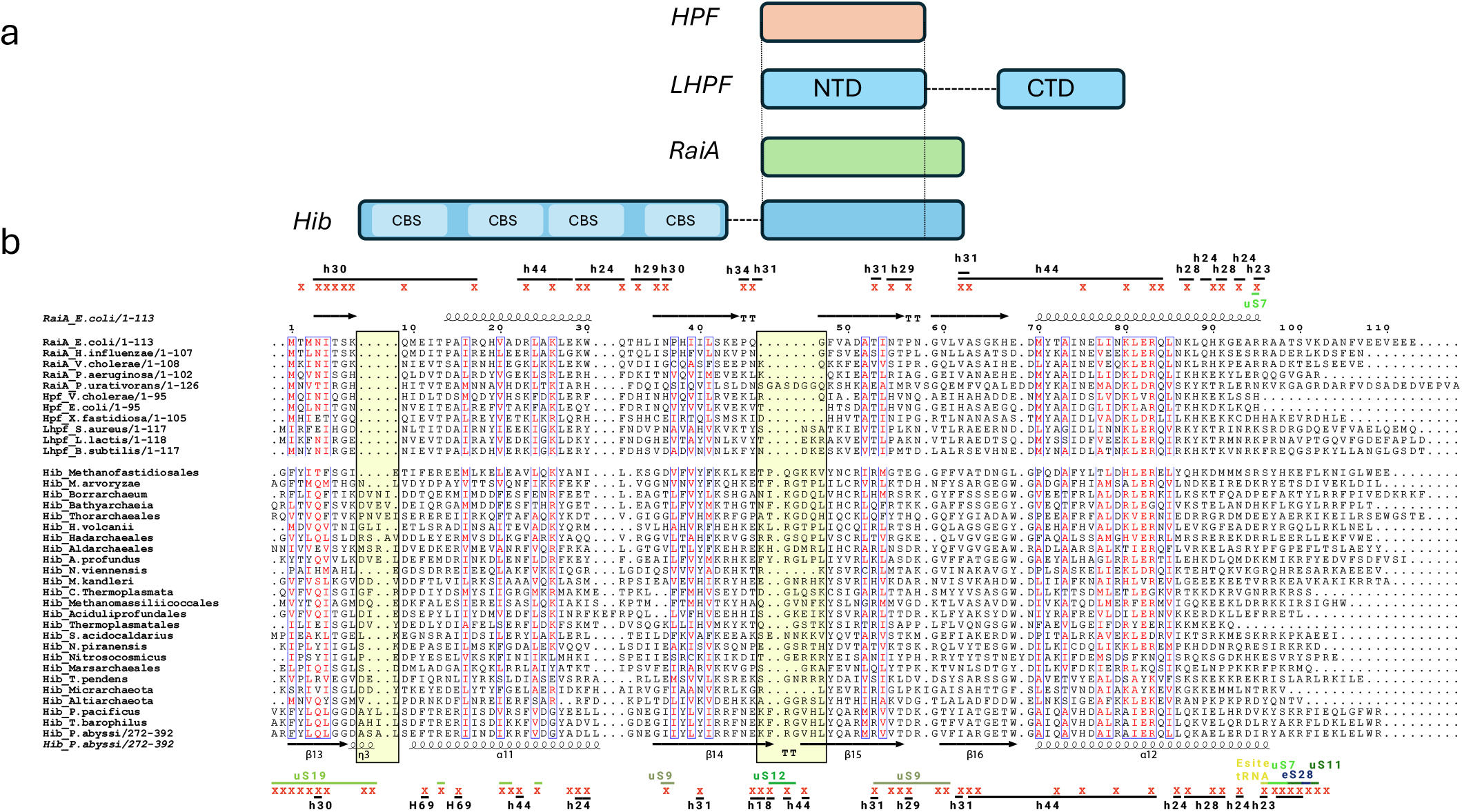
Hib shares a common domain with bacterial hibernation factors involved in 30S subunit binding. **a**. Schematic representation of domain arrangement of Hib and bacterial hibernation factors. Vertical dashed lines delineate homologous regions; horizontal dashed lines indicate variable length linker region between N- and C-terminal domains of LHPF and between N- and C-terminal domains of Hib. **b**. Structural alignment of C-terminal domains of Hib from diverse archaeal species with representative members of bacterial hibernation factor families. The reference structures are *E. coli* RaiA (4V8I) and *P. abyssi* Hib (present work). RaiA or Hib residues interacting with the rRNAs are indicated with dark dashes and labelled, those interacting with a ribosomal protein are indicated with colored dashes and labelled. The figure shows that most of the contact points of the hibernation factors with ribosomal RNA are conserved however *P. abyssi* Hib can be distinguished from RaiA homologues by two insertions (highlighted by yellow boxes). These insertions are involved in numerous contacts with ribosomal proteins (see text, Fig. 3 and ExtendedData 3). Note that RaiA from *Psychrobacter urativorans* has an insertion between the second and third b-strand. However, this region does not form a b-hairpin (PDB code 8RD8) as in Hib. H is used for helices of the LSU, h is used for helices of the SSU

**Supplementary Fig. 2:**
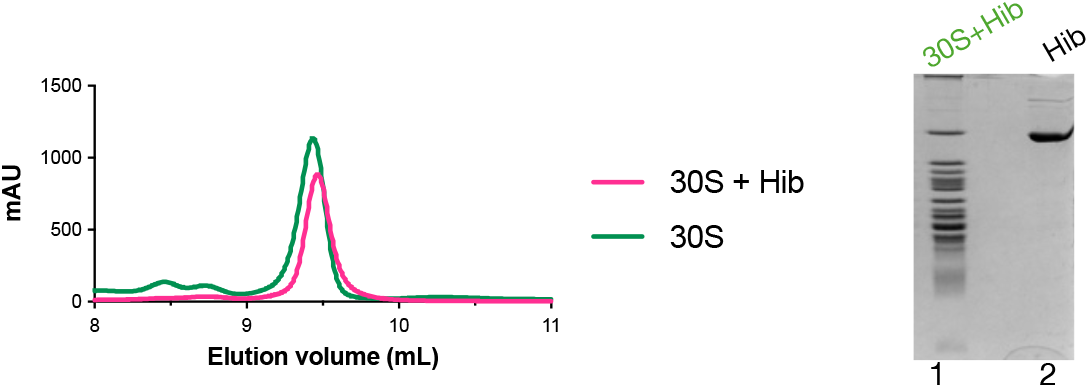
Bio-Agilent Sec5 chromatograms for P. abyssi ribosomal subunits, 30S (green), 30S+Hib mixture (pink). SDS-PAGE analysis is shown in the right-hand corner. 1: elution peak of the 30S+Hib mixture. 2: Hib. The analysis shows that Hib coelutes with the 30S fraction.

**Supplementary Fig. 3:**
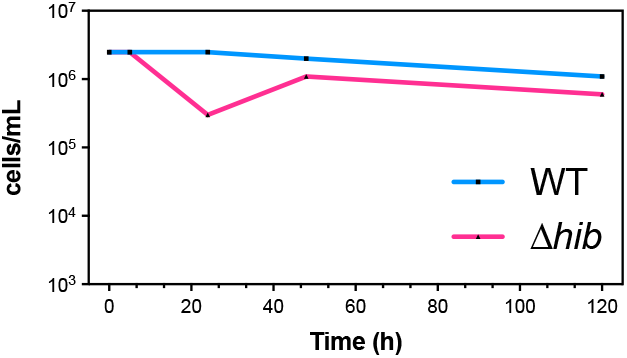
Viability after nutrient stress is unaffected by *hib* deletion in *T. barophilus*. Most Probable Number (MPN) assays were performed to assess the recovery of *T. barophilus* wild type (blue) and Δhib (orange) strains after incubation in carbon-free medium at 85 °C. Cell viability was monitored at different starvation times (0 h, 5 h, 24 h, 48 h, 120 h).

**Supplementary Fig. 4:**
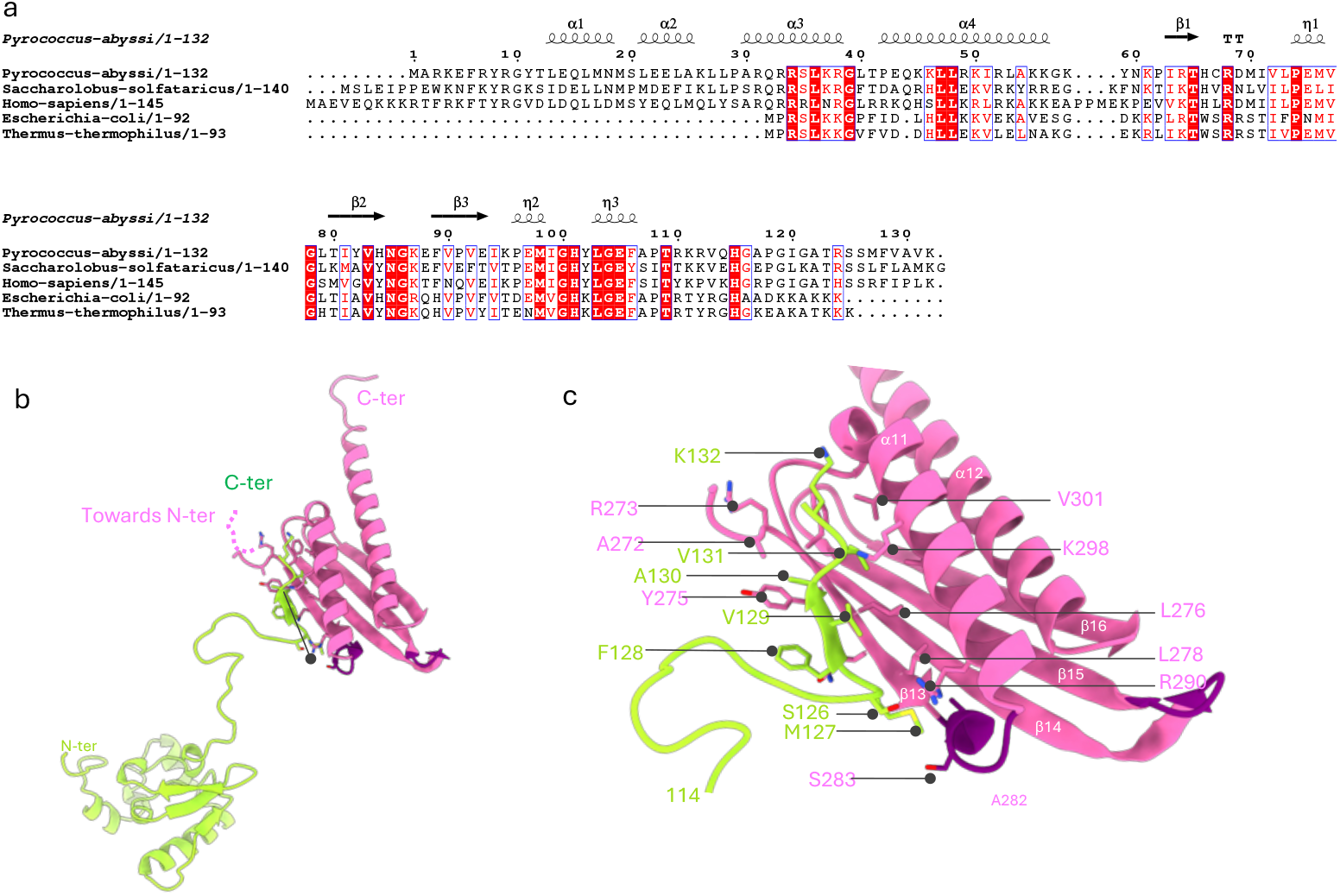
Interaction of the C-terminal domain of Hib with uS19. a. Sequence alignment of uS19 from the three domains of life. Eukaryotic and archaeal uS19 have a longer C-terminal tail (8 to 9 residues) than bacterial uS19, see also {Schmitt, 2020 #4351} and {Melnikov, 2018 #4250} for larger alignments. b. Interaction between uS19 and the C-terminal domain of Hib. The view highlights the contacts of the C-terminal tail of uS19 and the C-terminal domain of Hib. The Hib insertions, as compared to bacterial RaiA or HPF homologs are colored in dark violet. c. Close-up of the contacts between uS19 and the C-terminal domain of Hib.

**Supplementary Fig. 5:**
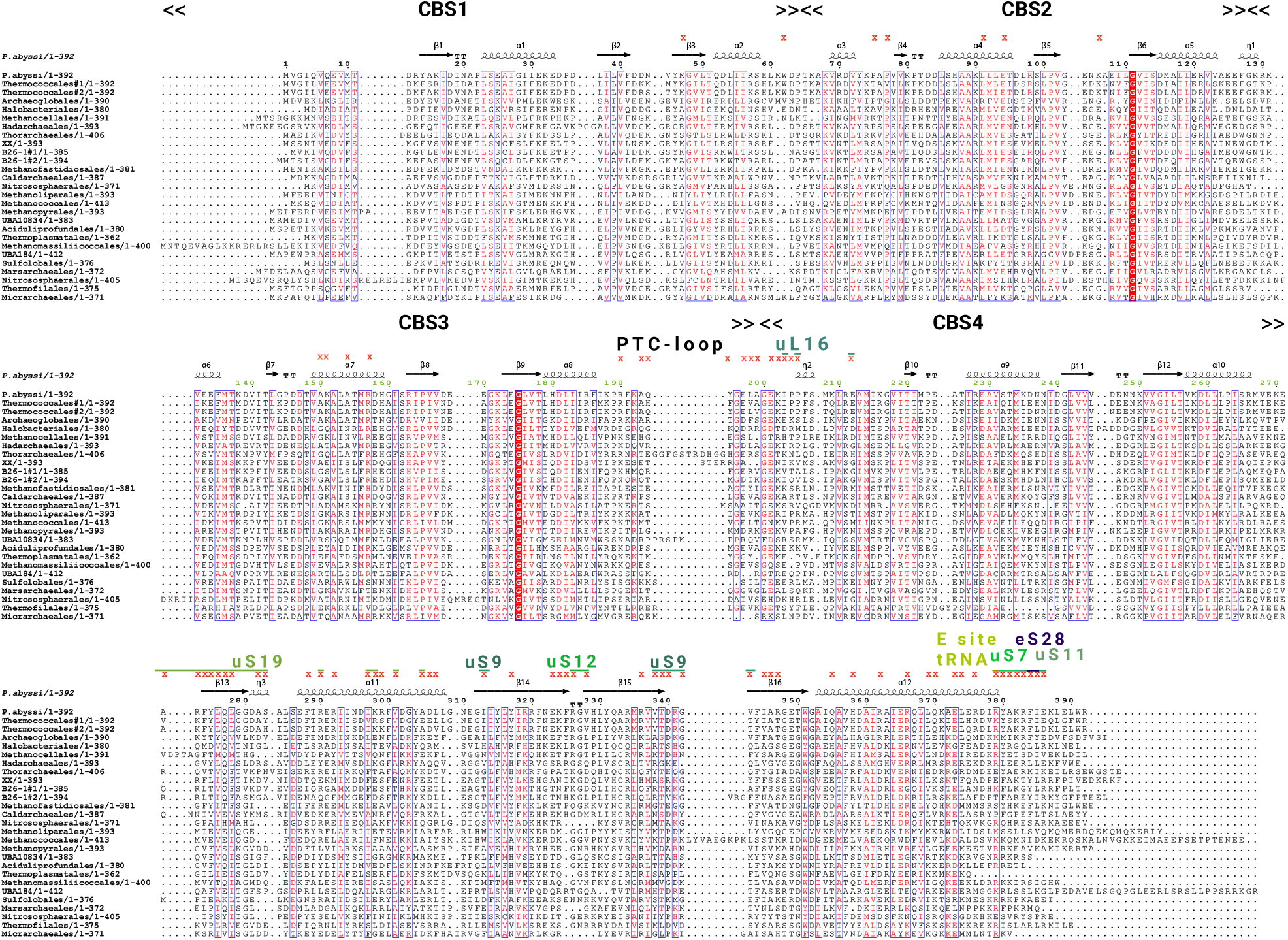
Sequence alignement of Hib from various Archaea. Hib residues interacting with the ribosome are indicated with a red star. When a ribosomal protein is involved in the interaction, it is indicated above the red star.

**Supplementary Fig. 6:**
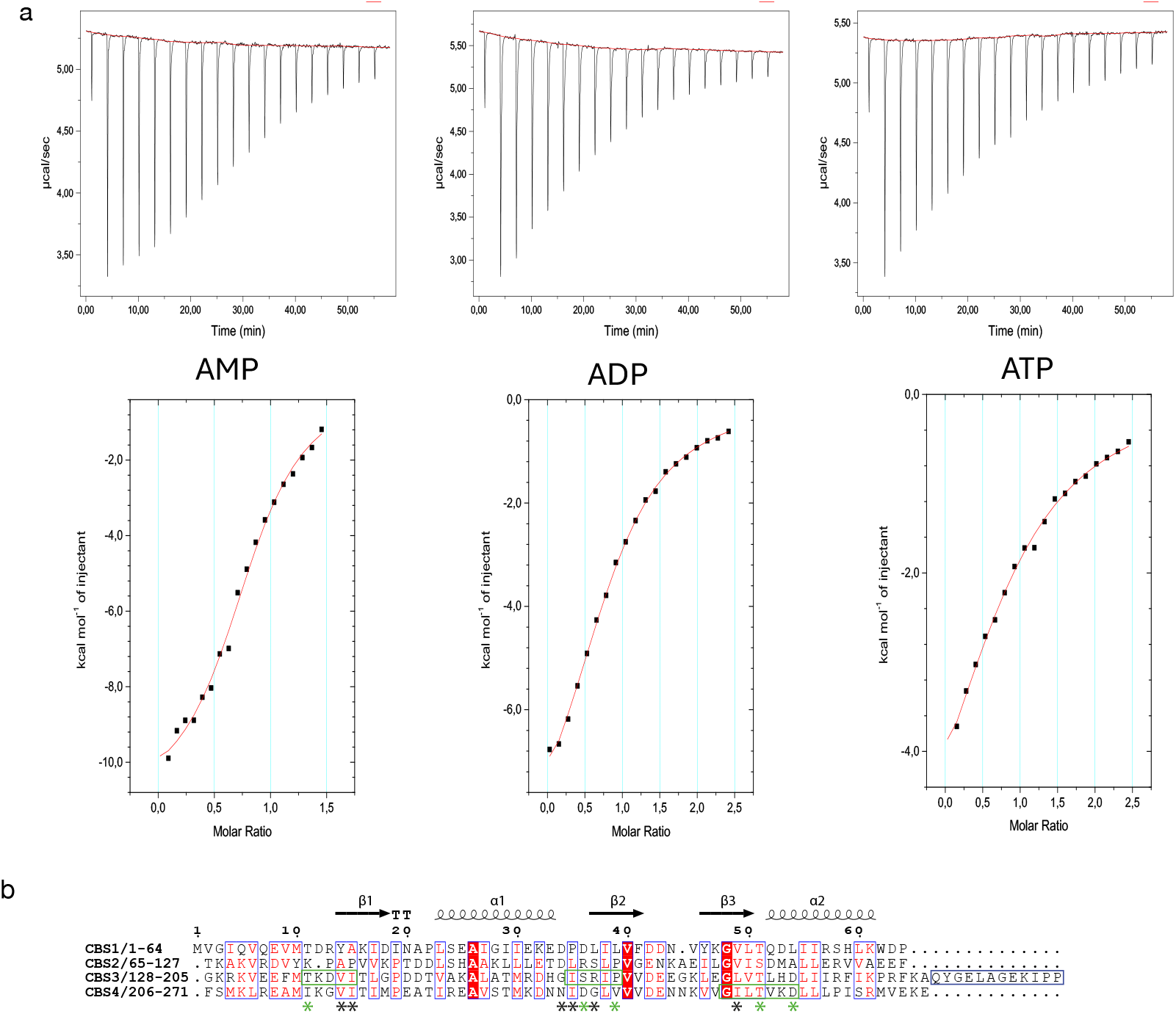
ATP binding. a. ITC titration curves (upper graphs) and corresponding binding isotherms (lower panels) for the interaction of Hib with AMP (left), ADP (middle) and ATP (right) in the presence of Mg^2+^ ions. Example experiments are shown. Derived Kd values and stoichiometries were 12 ± 2 µM (n=0.87 ± 0.05; AMP, 3 experiments), 65 ± 1 µM (n=0.82 ± 0.03; ADP, 2 experiments) and 113 ± 21 µM (n=0.81 ± 0.1; ATP, 3 experiments). b. Sequence alignement of the four Hib CBS domains. Conserved residues are framed in blue. ATP binds to the interface between CBS3 and 4. The residues involved in ATP binding are boxed in green and highlighted with stars. The green stars correspond to residues present in canonical binding sites of adenosine derivatives in a Bateman module {Ereño-Orbea, 2013 #4781}. Three blocks of residues are involved in ATP binding. The conserved second motif h-y-y’-h-P (where « h » is hydrophobic and « y » any residue) favors the interaction with adenosyl groups while preventing binding of guanosyl derivatives {Ereño-Orbea, 2013 #4781;Baykov, 2011 #4783}. Other residues, by their carbonyl groups favor the binding of adenosine derivatives as compared to guanosine derivatives. The third block of residues contains the G-h-h’-T/S-x-x’-D/N AMP binding motif. The aspartate residue of this motif is conserved and interacts with the 2’ and 3’ hydroxyl groups of ribose. Each CBS domain contains a potential binding cavity for nucleotides, yet a Bateman module typically binds only one adenylated ligand {Baykov, 2011 #4783}. Here, the absence of ATP binding in the second Bateman domain is explained by the presence of bulky side chains residues that obstruct the cavity and by the absence of conserved residues known to be important for adenosine derivatives binding. See also Fig. 5.

**Supplementary Fig. 7.**
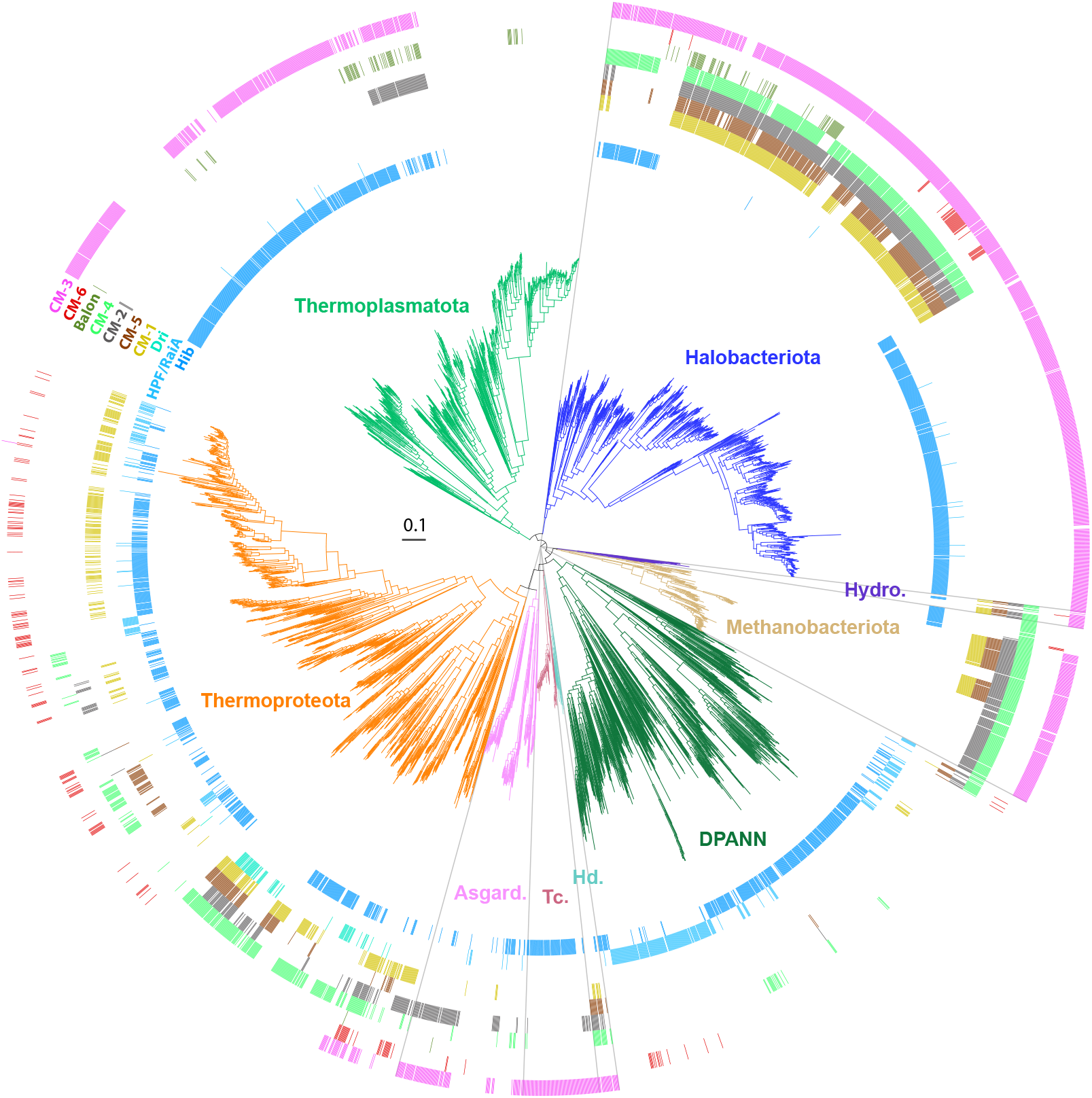
Mapping of hibernation factors and homologous CM proteins on a phylogeny of 4026 archaea. Hydro, Hydrothermarchaeota; Hd, Hadarchaeota; Tc, Thermococci; Asgard, Asgardarchaeota.

**Supplementary Fig. 8:**
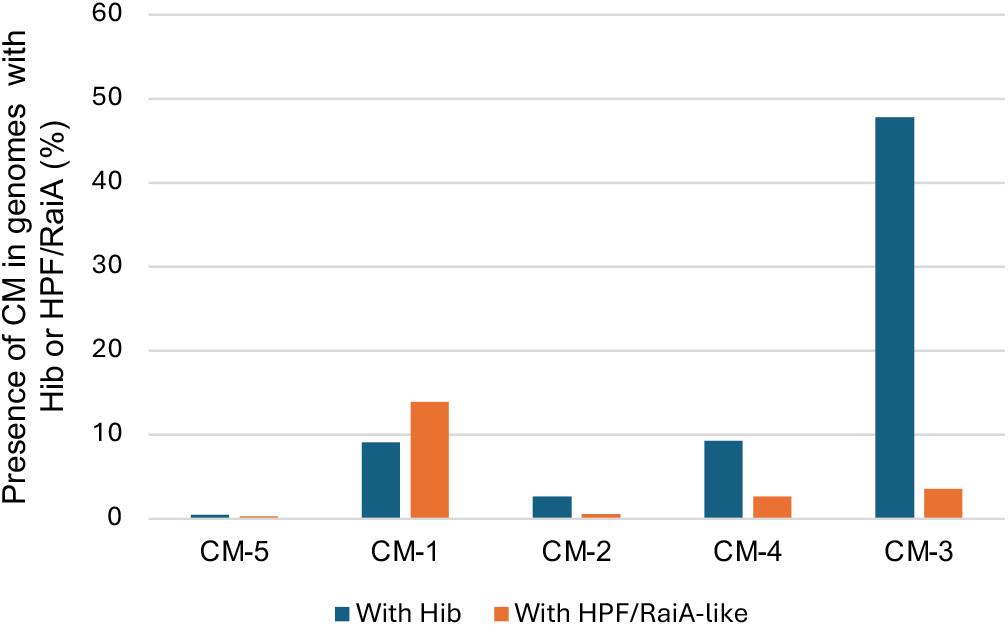
co-occurrence. Percentage of genomes coding for Hib or HPF/RaiA-like which also code for a standalone CBS module family (CM-5, CM-1, CM-2, CM-4, CM-3). Analysis done on a database of 4026 archaea.

**Supplementary Fig. 9:**
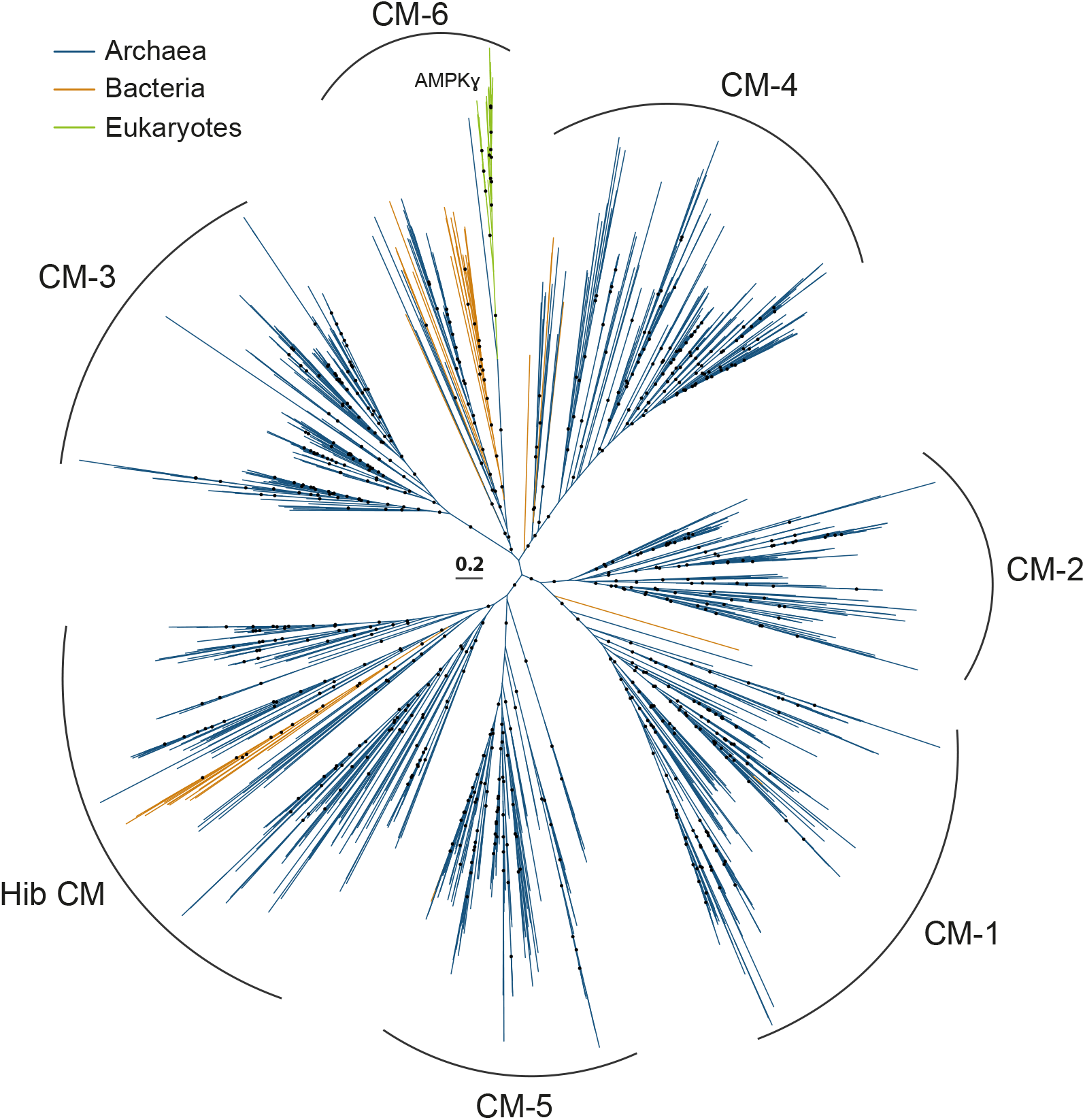
Phylogeny of CM. Tree of CBS modules (CM) including Hib CM, Dri and standalone CM from Archaea, Bacteria and Eukaryotes. AMPK1, AMP-activated protein kinase gamma subunit.

**Supplementary Fig. 10:**
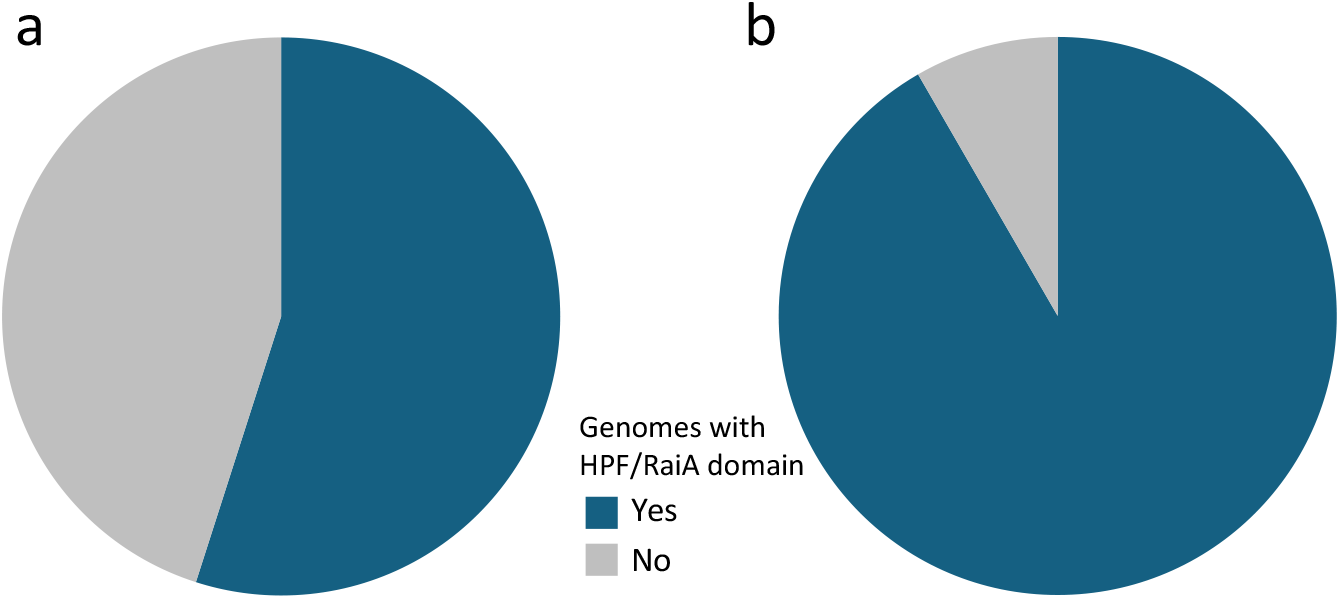
Distribution and diversity of proteins with a HPF/RaiA domain in archaea and bacteria. Proportion of **a**) archaeal and **b**) bacterial genomes coding for these proteins.

**Supplementary Fig. 11:**
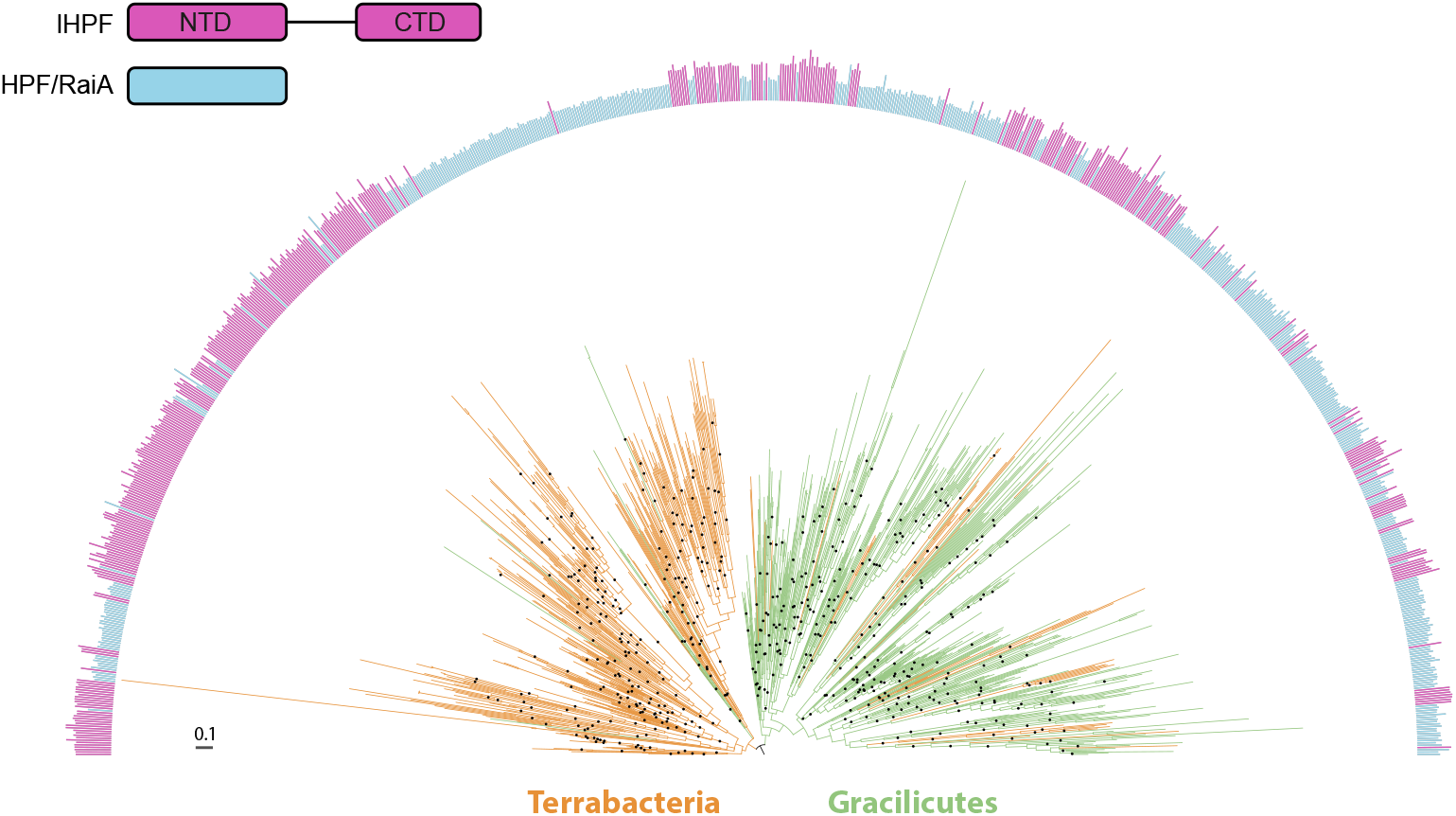
Phylogeny of the HPF/RaiA and lHPF of Bacteria. Classification between HPF/RaiA and lHPF categoies was not based on sequence length. HPF/RaiA were defined as sequences in which the CTD of lHPF(or another conserved domain) cannot be detected. Those in which this domain can be detected where defined as lHPF. Maximum-likelihood tree (LG+R7) based on a trimmed alignment of 85 amino acid positions. Black dotes on the branches indicate ultrafast bootstrap support >90%.

**Supplementary Fig. 12:**
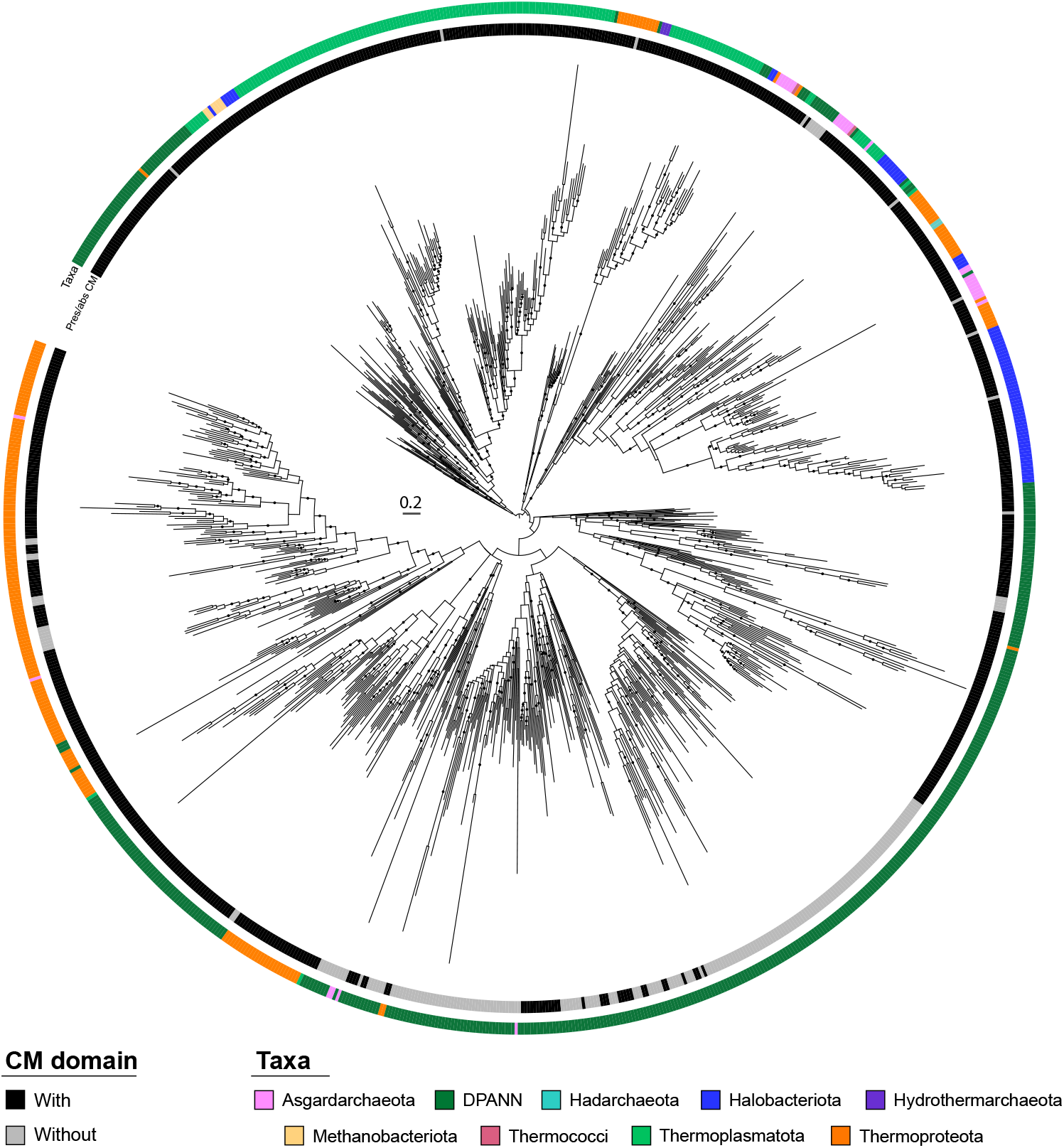
Phylogeny of the HPF/RaiA domain in Archaea. The first layer indicates the presence or absence of the CBS module (CM) in the respective proteins. The taxonomic affiliation of the sequences is indicated on the second layer. Sequences were clustered at 65% identity and a representative of each cluster was used for the phylogeny. Maximum-likelihood tree (LG+R5) based on a trimmed alignment of 97 amino acid positions. Black dotes on the branches indicate ultrafast bootstrap support >90%.

**Supplementary Fig. 13:**
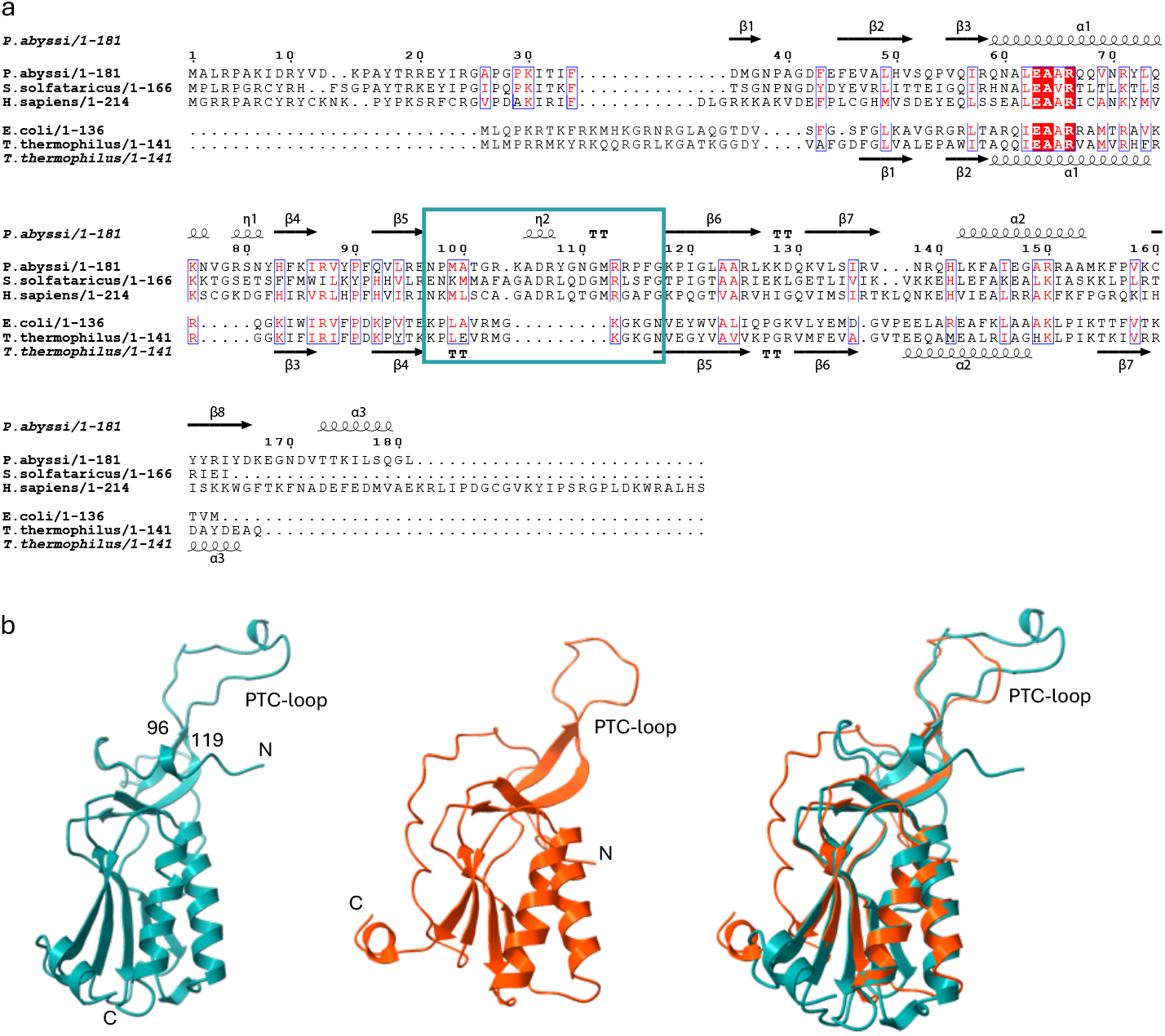
uL16 in the three domains of life. a. Sequence alignment of uL16 from the three domains of life. The uL16 loop located close to the PTC is squared. This loop is longer in eukaryotes and archaea and closer to the PTC. b. Comparison of *P. abyssi* uL16 (blue) with bacterial uL16 (T. thermophilus, red, PDB 4V8H).

**Supplementary Fig. 14:**
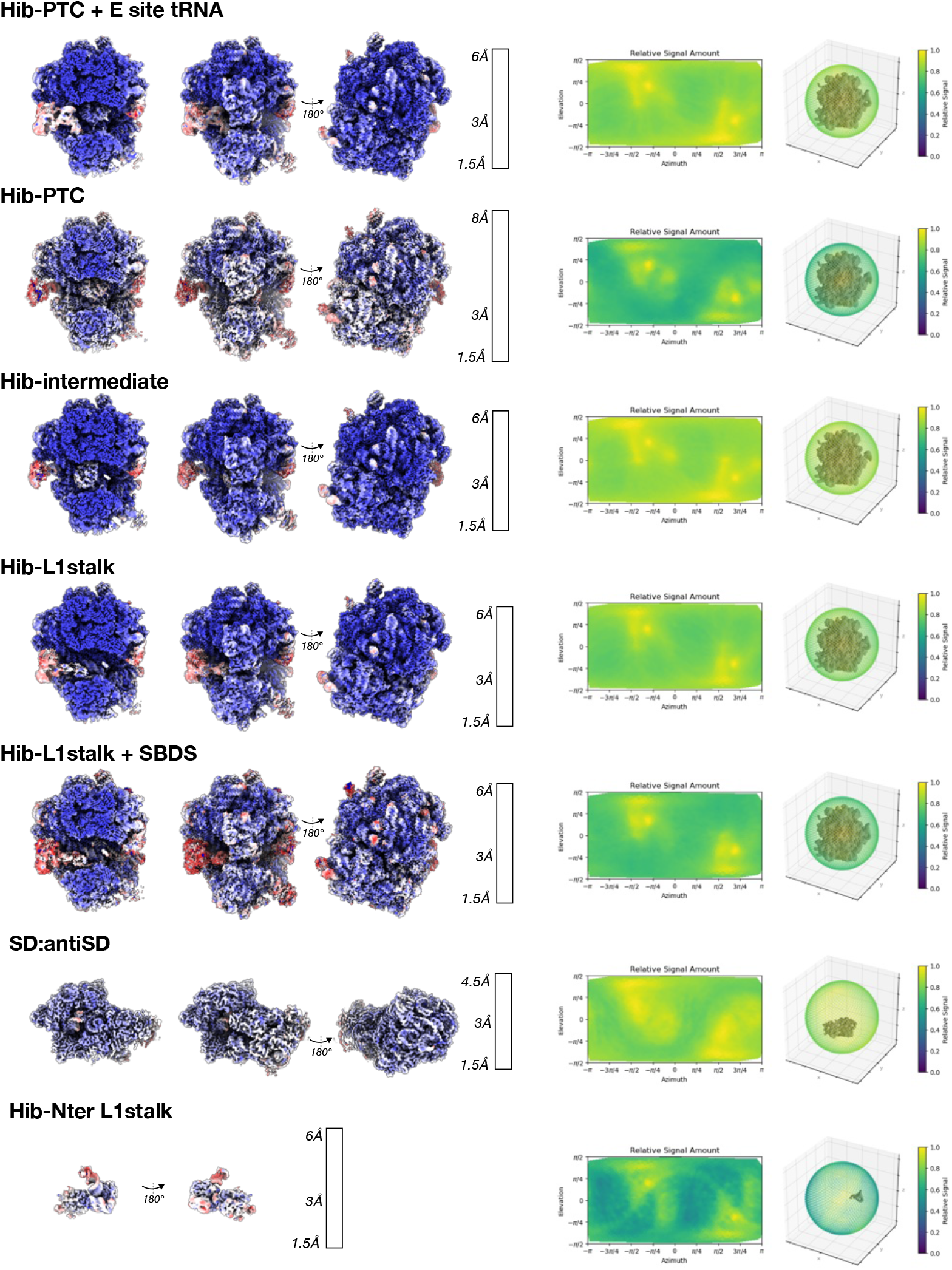
Local resolution and directional distribution of cryo-EM reconstructions. Local resolution for the different cryo-EM maps are shown on the left. Relative signal versus viewing direction, together with a 3D scatter plot are displayed on the right.

**Supplementary Fig. 15:**
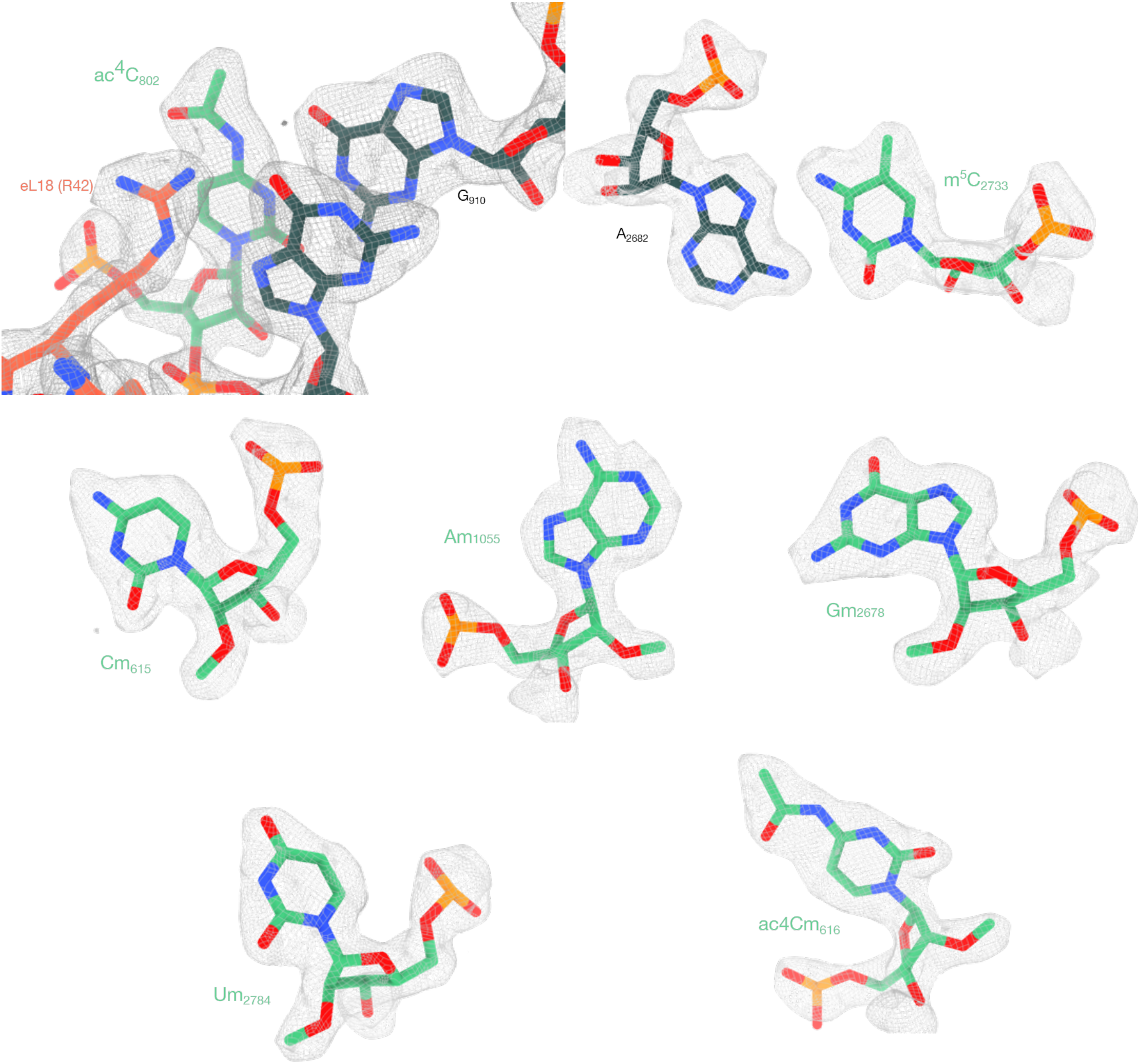
Examples showing the quality of the cryo-EM map for rRNA modified nucleotides and magnesium ions. The 2.1 Å resolution cryo-EM map (PDB 9SRE, EMDB 55139) is shown using the zone command in ChimeraX.

**Supplementary Fig. 16:**
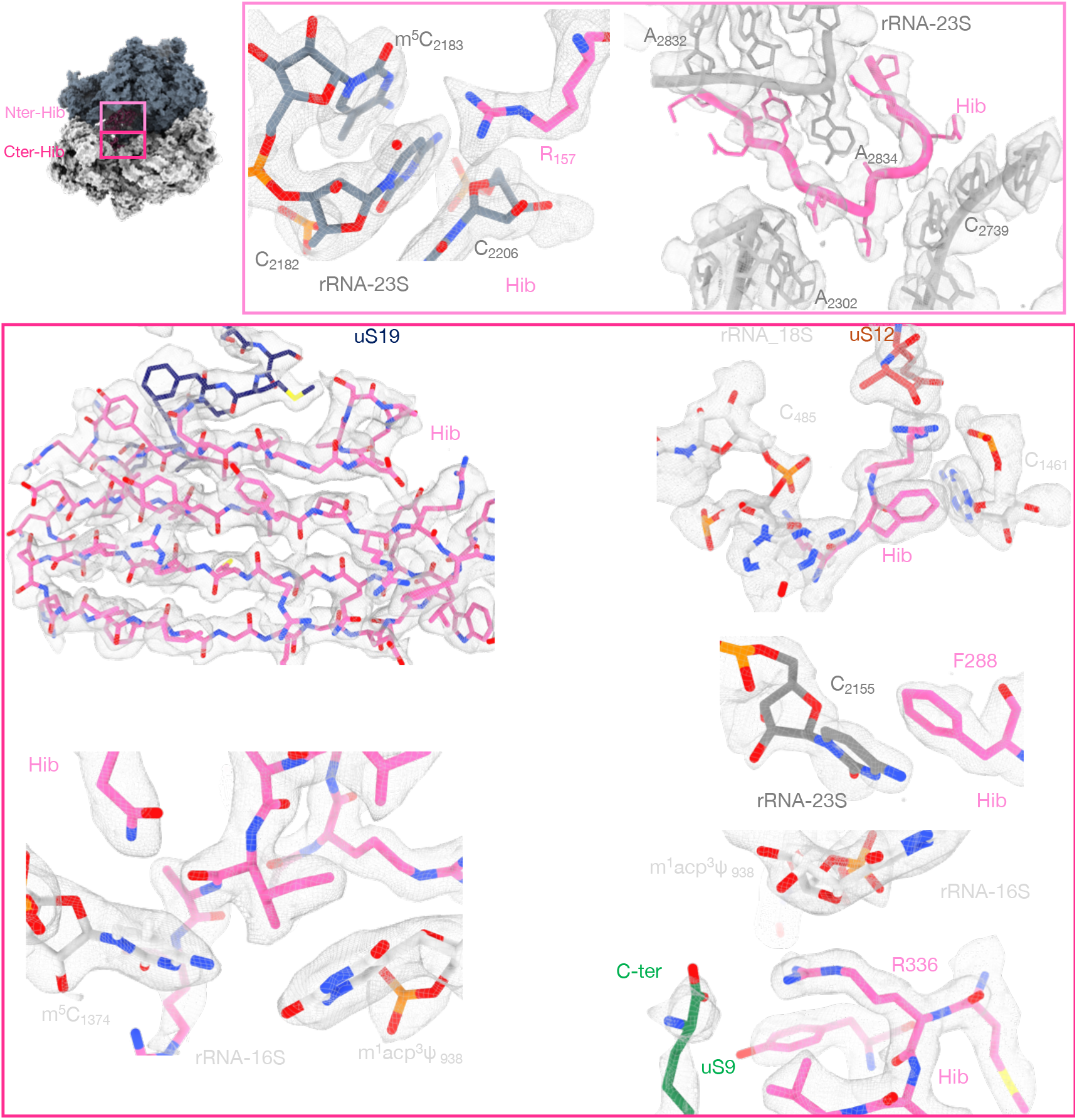
Examples showing the quality of the cryo-EM map around HIB protein. The 2.1 Å resolution cryo-EM map (PDB 9SRE, EMDB 55139) is shown using the zone command in ChimeraX.

**Supplementary Fig. 17:**
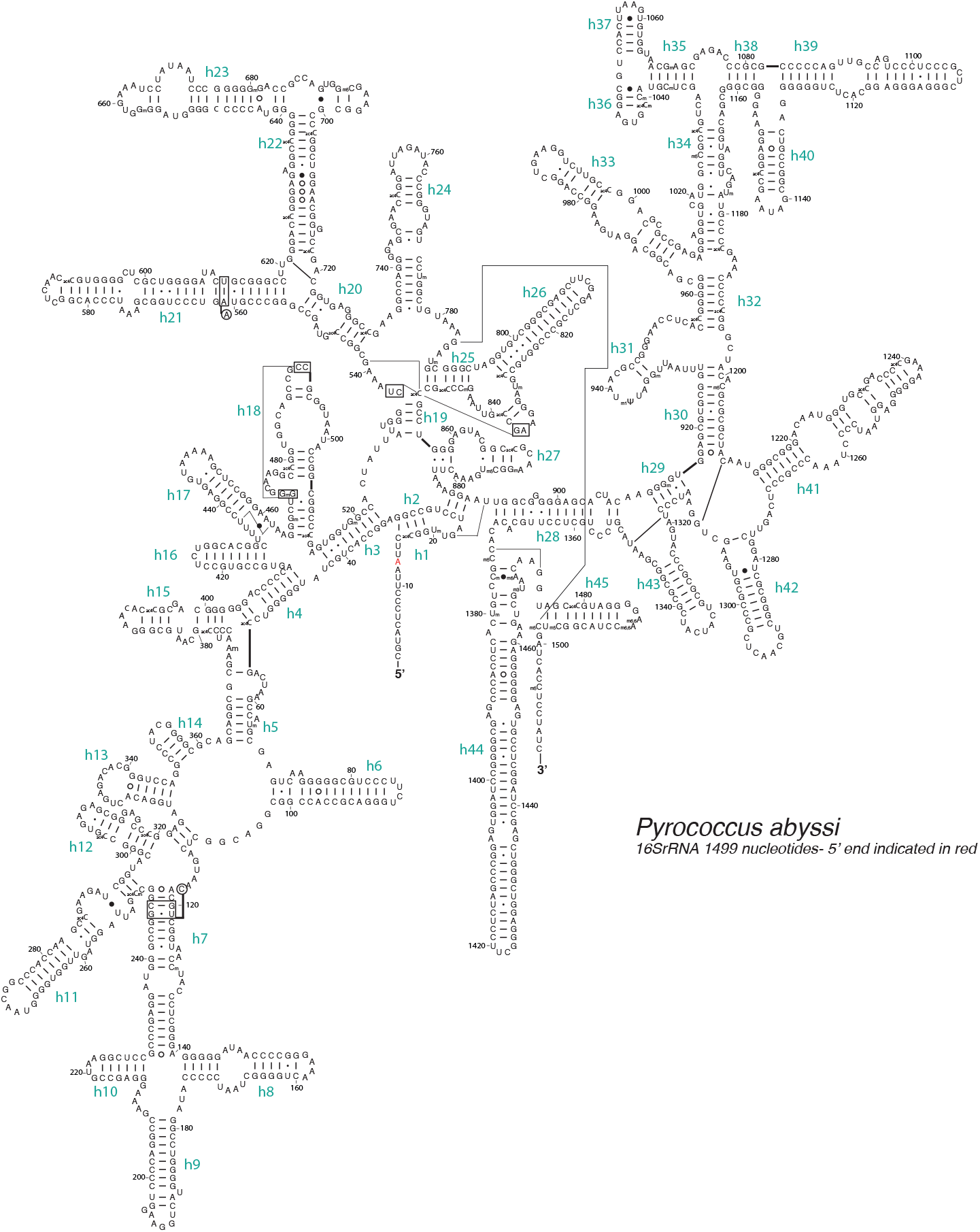
16S rRNAs from *P. abyssi*. Secondary structure diagrams were retrieved from https://crw-site.chemistry.gatech.edu/ and updated according to the cryo-EM structures. The 5’ end of *P. abyssi* 16S rRNA observed in the cryo-EM structure is indicated with a red letter {Bourgeois, 2025 #4769}.

**Supplementary Fig. 18:**
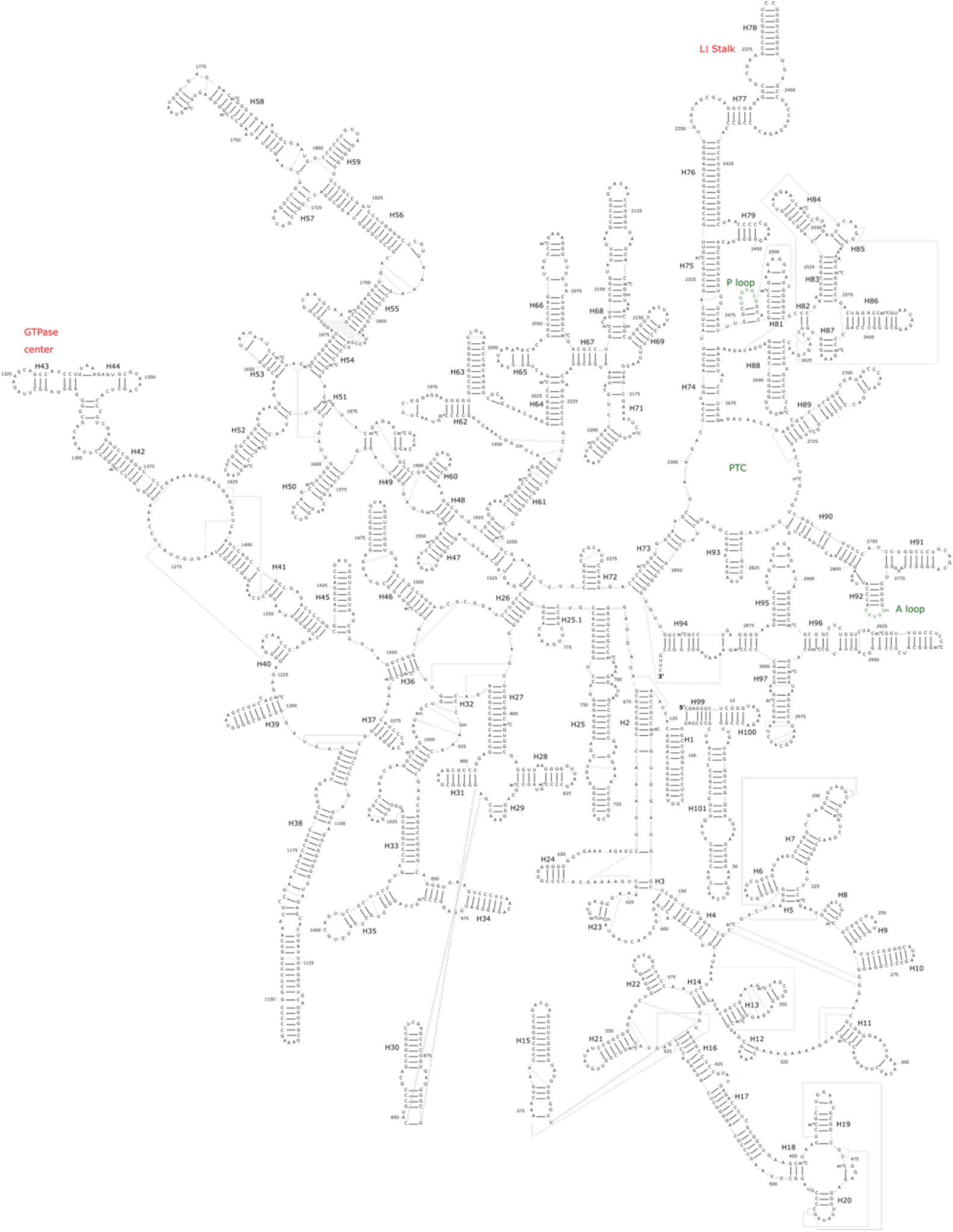
23S rRNAs from *P. abyssi*. Secondary structure diagrams were retrieved from https://crw-site.chemistry.gatech.edu/ and updated according to the cryo-EM structures. The 3’ and 5’ end of *P. abyssi* 23S rRNA observed in the cryo-EM structure are indicated.

